# Implications of noncoding regulatory functions in the development of insulinomas

**DOI:** 10.1101/2024.01.23.576802

**Authors:** Mireia Ramos-Rodríguez, Marc Subirana-Granés, Richard Norris, Valeria Sordi, Ángel Fernández Ruiz, Georgina Fuentes-Páez, Beatriz Pérez-González, Clara Berenguer Balaguer, Helena Raurell-Vila, Murad Chowdhury, Raquel Corripio, Stefano Partelli, Núria López-Bigas, Silvia Pellegrini, Eduard Montanya, Montserrat Nacher, Massimo Falconi, Ryan Layer, Meritxell Rovira, Abel Gonzalez-Perez, Lorenzo Piemonti, Lorenzo Pasquali

## Abstract

Insulinomas are rare neuroendocrine tumours arising from the pancreatic β-cells. While retaining the ability to produce insulin, insulinomas feature aberrant proliferation and altered hormone secretion resulting in failure to maintain glucose homeostasis.

With the aim of uncovering the role of noncoding regulatory regions and their aberrations to the development of these tumors, we coupled epigenetic and gene expression profiling with whole-genome sequencing. As a result, we mapped H3K27ac sites in the tumoral tissue and unraveled overlapping somatic mutations associated with changes in regulatory functions. Critically, these regions impact insulin secretion, tumor development and epigenetic modifying genes, including key components of the polycomb complex. Chromatin remodeling is apparent as insulinoma-selective regions are mostly clustered in regulatory domains, shared across patients and containing a specific set of regulatory sequences dominated by the binding motif of the transcription factor SOX17. Moreover, a large fraction of these regions are H3K27me3-repressed in unaffected β-cells, suggesting that tumoral transition is coupled with derepression of β-cell polycomb-targeted domains.

Our work provides a compendium of aberrant *cis*-regulatory elements and transcription factors that alter β-cell function and fate in their progression to pancreatic neuroendocrine tumors and a framework to identify coding and noncoding driver mutations.

## Introduction

The pancreas is an heterogeneous tissue hosting some of the most debilitating diseases, including diabetes mellitus, and cancer of the exocrine and the endocrine tissue compartments^1^. About 35% of pancreatic neuroendocrine tumors (PNETs) are hormone-secreting (also defined as functional PNETs), with insulinoma being the most prevalent among them. Insulinomas are slow-growing adenomas derived from the β-cells that constantly produce insulin or proinsulin^2–4^. They are rare, occurring in only 4 individuals per million each year. The identification of insulinomas in medical settings is typically triggered by the excessive production of insulin, leading to hypoglycemia and associated psychomotor symptoms. Due to their rarity, they are not included in comprehensive cancer genomic surveys such as The Cancer Genome Atlas (TGCA) or the International Cancer Genome Consortium (ICGC). Although about 10% of them carry germline or somatic mutations of the *MEN1* gene, and several groups recently reported recurrent mutation affecting the transcription factor (TF) YY1^5,6^, the mechanisms underlying β-cell overgrowth and neoplastic transformation are still obscure. Several studies point to the possible involvement of both genetic and epigenetic mechanisms in the tumour development and loss of β-cell identity^3,7,8^, yet the noncoding regulatory landscape and the full genetic profile of these tumors has not been yet elucidated.

The human genome sequence contains the instructions to generate a vast number of cell fate programs. This is possible because each cellular state utilizes distinct sets of genomic regulatory regions. Once reached their final differentiation stages, adult cell fate is actively maintained by reinforcement of specific regulatory networks, encoded in chromatin states, defined in part by the complement of active *cis*-elements. An increasing number of studies have demonstrated that epigenomic reprogramming, especially enhancer reprogramming, plays an important role in cancer progression and metastasis^9^. Similarly, TFs have crucial roles as agents driving and adjusting the reprogramming process, and have been described to initiate oncogenic processes by activating well-defined functions^10^.

A central feature of tumor development is the acquisition of somatic mutations. Large cancer genomic studies including the Pan-Cancer Analysis of Whole Genomes (PCAWG) show that a large fraction of all somatic mutations lie in non-protein coding DNA regions, including variants overlapping known regulatory annotations. These observations suggest that alteration of noncoding functions could underlie driver events in the acquisition of a cancer phenotype^11^. Nonetheless, currently there is a lack of understanding of the role of the noncoding genome in cancer^12,13^, limiting our overall understanding of the regulatory programmes intervening and driving tumoral cell states.

We have now profiled transcriptional maps, *cis-*regulatory networks and genome wide annotations of somatic genetic aberrations in insulinomas. We exploit these data to uncover aberrant regulatory functions defining the tumoral state. These analyses permit elucidation of the functional mechanisms driving the β-cell neoplastic transformation.

## Results

### Mapping the regulatory landscape of insulinomas

We profiled the transcriptome of a total of 17 insulinoma samples, including 6 obtained from a prior report^3^ (**Figures 1A**, **S1A** and **Table S1**). We next compared them with those of unaffected human pancreatic islets^14–17^ and insulin-producing β cells^17–20^ (**Figure S1A**) and found that insulinomas cluster together and separately from the normal tissues regardless of isolation technique or center of origin (**Figure S1B-C**). Differential expression analysis uncovered ∼1,100 genes upregulated in the tumor samples (**Figures 1B** and **S1D,** and **Table S2**). In line with previous reports^3^, we found that these genes are enriched in chromatin regulators and modifiers (**Figures 1C** and **S1E**), and histone acetyltransferases in particular (**Figure S1F**). Driven by this observation, we sought to explore whether the tumor development is associated with reshaping of the regulatory landscape of β-cells. We used chromatin immunoprecipitation coupled with next generation sequencing (ChIP-seq) to profile H3K27ac on 12 insulinoma samples, 11 of which had matching RNA-seq data, to map active regulatory elements (RE), including transcriptional promoters and enhancers (**Figure 1A** and **Table S1**).

**Figure 1:**
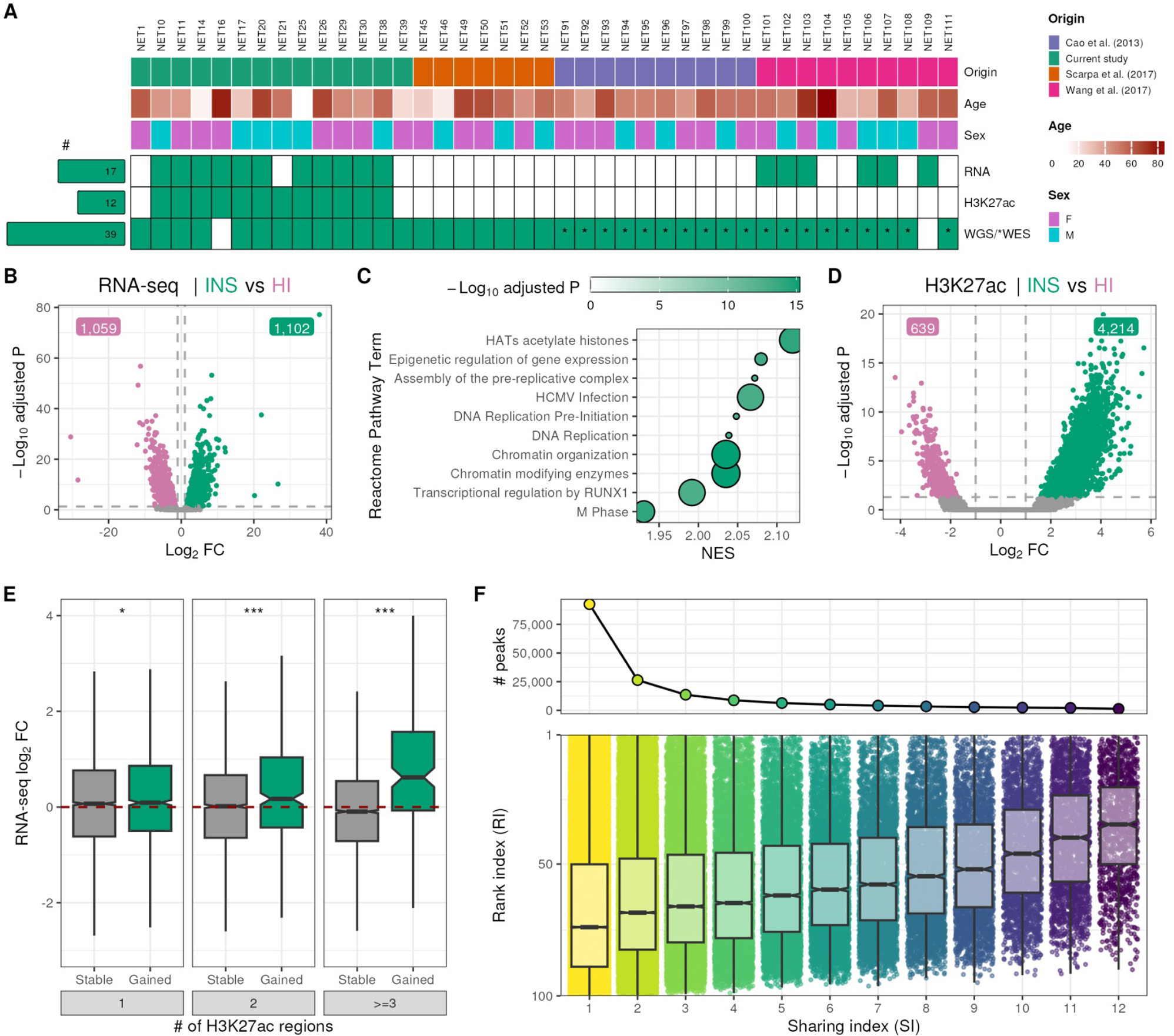
Mapping the transcriptome and regulatory landscape of insulinomas. **A**, Summary of insulinoma samples and assays included in this study. Top rows show the sample origin and patient age and sex. Volcano plot of differentially expressed genes (**B**) and H3K27ac enriched regions (**D**) in insulinomas (green) compared to untransformed human pancreatic islets (HI, pink). Dotted lines show thresholds for significance (|log2 fold change| > 1 and adjusted P < 0.05). Green: up-regulated genes or gains in H3K27ac; Pink: down-regulated genes or losses in H3K27ac. Grey: stable genes or H3K27ac regions. **C**, GSEA of Reactome Pathway terms enriched in insulinoma-selective genes (as compared to HI) are strongly dominated by histone modifier enzymes and chromatin remodelers. **E**, Distribution of gene expression changes in insulinoma vs. HI for transcripts in the vicinity of increasing numbers of H3K27ac regions shared with the normal tissue (stable) or insulinoma-selective (gained). Two-sided Wilcoxon test: * P < 0.05, *** P < 0.001. **F**, Sharing and rank indexes derived from H3K27ac signal in insulinomas in regions distal to gene TSS.

Overall we mapped a total of 12,454 proximal promoters and 49,259 distal putative enhancers across all insulinoma and control human islet samples^19,21–23^, resulting in a comprehensive map of active REs (**Figure S2A-B**), capable of differentiating accurately between insulinomas and the control tissues (**Figure S2C-D**). Next we identified ∼5,000 differential H3K27ac enrichments in insulinomas compared to untransformed human islets (**Figure 1D** and **Table S3**). We observed that insulinoma-selective H3K27ac sites are mostly distal to transcriptional start sites (TSS) (7% proximal and 93% distal, **Figure S2E**). Remarkably, we found that gains in H3K27ac enrichment are linked to upregulation of the nearby gene/s. Moreover, these changes are highly correlated with the number of associated H3K27ac sites, suggesting a cumulative effect of the regulatory elements on the expression of the nearby transcripts (**Figure 1E**).

Genetic and epigenetic heterogeneity is a hallmark of cancer. Unsupervised clustering revealed that genome-wide enrichment of H3K27ac can clearly distinguish insulinoma from pancreatic islets and non-functional PNETs, as well as from other cancer types^24–29^ and untransformed cell types^19,21–23,27,28^ (**Figure S3A**). Yet, insulinoma’s transcriptional program remains closer to their cells of origin when compared to other cell types (**Figures S1B-C** and **S3A**). These results, as well as the comparatively low inter-patient variability of the H3K27ac signal (**Figure S3B),** suggest lower heterogeneity of the regulatory functions in insulinoma, as compared to non-functional PNETs and other cancer types. We nevertheless sought to infer whether the H3K27ac epigenetic hallmark is shared between different patient samples (inter-sample heterogeneity) and representative of the major sample cell clones (intra-sample heterogeneity). To this end, we computed a sharing index (SI) by annotating the number of patients sharing each H3K27ac enriched site and next ranked these regions by their signal intensity, thus deriving the Rank Index (RI)^26^. The rationale of the RI metrics stands on the observation that heterogeneity within the cell population was demonstrated to be the major contributor to H3K27ac signal intensity^26^. In our insulinomas cohort we observed a strong correlation between SI and RI, indicating that, as for other tumours^26^, clonal epigenetic events are those that are more shared between different patients (**Figures 1F** and **S3C**). Moreover, we found that 50% of insulinoma-selective sites were common to more than 67% patient samples (**Figure S3D**). Altogether these findings suggest that insulinomas from different patients may share common mechanisms of gene expression deregulation.

Overall, we mapped a first draft of active regulatory regions relevant to tumorigenesis in insulinoma. Our data suggest that the genome-wide aberrant deposition of H3K27ac in these tumors tends to be related to gene expression regulatory functions and shared between patients.

### Recurrent coding mutations in insulinoma are rare

In order to assess the contribution of genetic alteration to the neoplastic transformation in insulinomas, we sequenced the whole genome (WGS) of 13 tumors and patient-matched peripheral blood cells. We next integrated the newly generated data with published WGS^30^ and whole exome sequencing (WES)^3,5^ to obtain a large dataset (n=40) of paired tumor-normal samples, for 10 of which we had matching RNA-seq and H3K27ac data (**Figure 1A**). We focused on samples not carrying germline mutations in *MEN1*, a known insulinoma driver gene^31^, in order to facilitate the discovery of yet undescribed driver mutations.

For the detection of somatic mutations in insulinomas, we employed a standardized set of established algorithms for alignment and variant calling (**Figure S4A**) and implemented rigorous variant filtering procedures. We unveiled 24,627 single-nucleotide variants (SNVs) and 870 small INDELs (626.45±668.70 SNVs and 11.97±14.12 INDELs per tumor) (**Figure S4B**). Globally, we observed a low mutation burden (median: 0.37 mutations per megabase; range 0.06–1.98) as compared to a large panel of tumors ^32^ (n=33), including pancreatic tumors arising from the exocrine tissue (pancreatic ductal adenocarcinoma; median: 0.98 mutations per megabase; range 0.03–389.27) (**Figure S4C**).

We next used SigProfiler^33^ to decompose *de novo* extracted signatures and match them to the reference COSMIC signatures^34^. Three single base substitution clock-like signatures (SBS) related to aging (SBS1, SBS5 and SBS40, with cosine similarity of 0.98) constituted the first-described mutational footprint of insulinomas (**Figure S4D**). Interestingly, similar mutational signature patterns have been previously described in pancreatic ductal adenocarcinoma (PDAC), suggesting that only endogenous processes likely contribute to both types of pancreatic tumors^35^.

We annotated a total of 1,064 somatic mutations (1,063 SNVs and 1 INDEL, 26.6±22.5 variants per sample) to genomic coding sequences (**Table S4**). With the exception of the previously described *YY1* T372R mutation^5,6^, which appeared in 17% of the patients in our cohort, and in line with previous reports^3^, recurrent coding mutations in insulinoma were rare. Nevertheless, in multiple patient samples, we observed recurrent mutations in several genes, namely *BRD1*, *CFAP47*, *COL11A1*, *ZZEF1*, and *RNF213*, which had not been previously associated with the development of this tumor. Moreover, *RNF213*, an E3 ubiquitin ligase protein, was identified as a potential driver gene through a pipeline combining seven state-of-the-art computational methods to identify genes under positive selection across tumors^36^ (**Figure 2A**).

**Figure 2:**
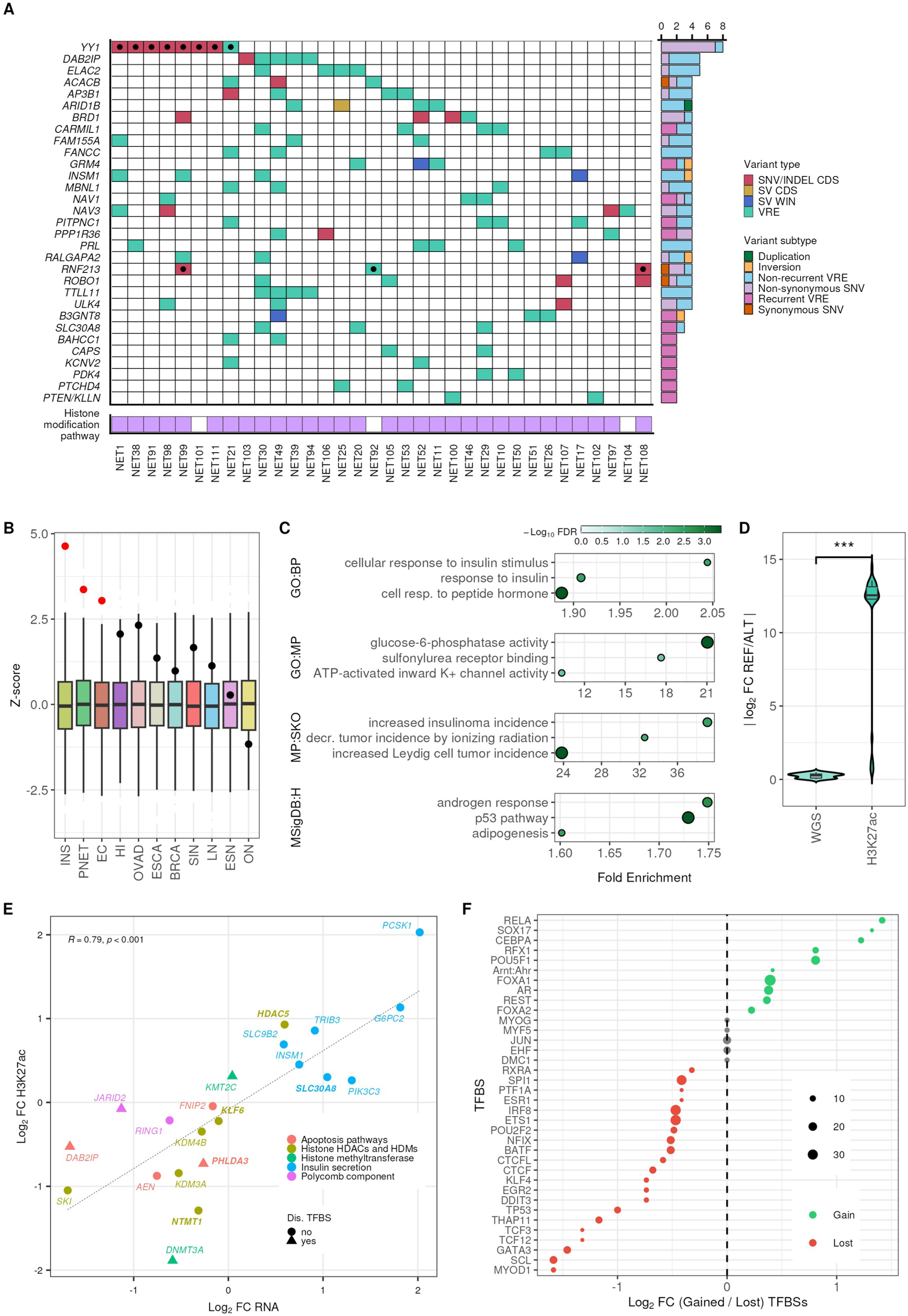
Genome-wide mutation landscape in insulinomas. **A**, Ranking list of the genes accumulating the highest frequency of mutated genomic elements, including coding exons and nearby mutated H3K27ac enriched sites (VREs), observed in a cohort of 40 insulinoma tumours. Only somatic variants are included. Right, the number of samples affected by any of the genomic alterations in each gene and the category. Black dot denotes coding driver genes inferred by IntOGen^36^. Bottom, samples affected by any genomic alteration in at least one gene related to the histone modification pathway (GO:0016570). **B**, Variant set enrichment analysis illustrating that insulinoma somatic mutations are enriched at H3K27ac sites active in insulinoma and its cell of origin (β-cells), as well as in non-functional PNETs. However, no overreppresentation of these variants was observed in H3K27ac sites active in other types of cancer or untransformed primary tissues. Significant enrichment scores are shown in red (Benjamini–Hochberg-adjusted P <0.05). The boxplot limits show the upper and lower quartiles; the whiskers extend to 1.5× the interquartile range; the notch represents the median confidence interval for distributions of matched null sets (500 permutations). INS = Insulinoma, PNET = Non-functional Pancreatic Neuroendocrine Tumour, EC = EndoC-βH1, HI = Human pancreatic Islets, OVAD = Ovarian Adenocarcinoma, BRCA = Breast Carcinoma, ESCA = Esophageal Squamous Cell Carcinoma, BRCA = Breast Carcinoma, LN = Lung Normal, ON = Ovarian Normal and SIN Small Intestine normal **C**, Genes associated with VREs are implicated in β cell function and neoplastic transition. Gene Ontology: Biological Process (GO:BP), Gene Ontology: Molecular Function (GO:MF), Mouse Phenotype Single KO (MP:SKO) and Human Molecular Signatures Database: Hallmark (MsigDB:H). **D**, Violin plots showing the absolute allele fold change distribution of WGS and H3K27ac ChIP-seq reads, carrying (ALT) or not (REF) the mutated genotype at VREs. The data show a significant allelic skew of H3K27ac reads, indicating that the histone modification deposition at VREs depends on the somatic mutation genotype. The analysis is based on 30 VREs for which both WGS and H3K27ac matched data were available. Two-sided Wilcoxon test ***P < 0.001. **E**, H3K27ac and gene expression fold change in insulinoma (Insulinoma mutated vs. insulinoma wildtype) at selected VREs and their associated transcript, A triangle indicates that the mutation/s is/are predicted to affect the sequence by creating or disrupting a TF binding site. Color denotes the pathway in which the target gene is implicated. Genes associated to multiple VREs are depicted in bold. **F**, List of TF binding sequences (TFBS) predicted to be created (gained) or disrupted (lost) by somatic mutations at VREs. The size of the circle is proportional to the number of modified TFBS, while the color depicts a higher ratio of created (green, eg. SOX17) or disrupted (red, eg. TP53) binding sequences.

To build an accurate and comprehensive somatic structural variants (SV) truth set, we used a combinatorial approach in which we retained consistent results obtained from four independent SV algorithms callers (See Methods). To minimize the detection of false-positives alterations, we applied 1) stringent quality control and removal of known population SV and 2) visual validation^37^ (**Figure S4A**). We recovered a total of 146 SV the majority of which (55.5%) arose from chromosomes 6, 7 and 12 (**Figure S4E** and **Table S4**). Interestingly, within the transcripts affected we annotated histone modifiers and genes related to the polycomb complex (*EZH2*, *HDAC2* and *KMT5A*), as well as others already known to be involved in insulinoma development (*INSM1* and *PTPRN2*) (**Figures 2A**, **S4F** and **Table S4**)

### The impact of noncoding somatic mutations

The noncoding genome is populated by functional regulatory elements playing a critical role in regulating gene expression and in maintaining a cell-type specific phenotype. We thus addressed whether somatic regulatory variants affecting non-coding regulatory elements are implicated in driving the tumoral phenotype.

By mapping the identified somatic mutations to the newly generated regulatory maps in insulinoma we found an overrepresentation of mutations at H3K27ac sites (adjusted P<0.02, *z*=4.64, **Figure 2B**). Similarly the mutations were enriched at H3K27ac sites active in normal β cells and PNETs. It’s worth noting that the genomic distribution of the mutations may be driven by the tissue-specific chromatin landscape of the tumoral cell type of origin^38,39^, and may not represent the result of a tumoral-driven positive selection process. However, this finding presents an opportunity to explore how somatic mutations in insulinoma may impact β-cell tissue-specific regulatory functions as well as pathways involved in neoplastic processes.

We thus define as Variant Regulatory Element (VRE) an insulinoma or human islet H3K27ac site bearing an insulinoma somatic mutation (**Table S5**). Several observations suggest a functional role of VREs: 1) VREs are preferentially located proximal to gene transcription start sites (TSS) (∼40% are located <2Kb from a TSS, P<7.49×10^71^ (**Figure S5A**), 2) their sequence is, on average, more evolutionarily conserved as compared to matched control H3K27ac sites (**Figure S5B**) and 3) they are enriched for specific TF binding sites including Insulin gene enhancer protein (ISL-1), MAF BZIP Transcription Factor A (MAFA), Forkhead Box O1 (FOXO1) and SMAD Family Member 3 (SMAD3), a factor related to canonical signaling cascade of TGF-β and previously described to have a role in the development of insulinomas^3^ (**Figure S5C**). Moreover, 4) VREs are located at the promoter or in physical proximity to genes clearly implicated with insulinoma and tumoral developmental functions, including genes already known to be implicated in insulinoma progression (P=1.26×10^−3^), insulin response and secretion (P=5.20×10^−6^), glucose-6-phosphatase activity (P=4.67×10^−5^) and the p53 pathway (P=4.70×10^−4^) (**Figure 2C**).

To gain insight into the potential functional role of noncoding mutations on regulatory genomic functions, we took advantage of samples with matched WGS/ChIP-seq/RNA-seq to assess the relationship between genotype and both local enrichment of H3K27ac in VREs and changes in expression of the nearby genes. At VREs, we found a significant allelic skew for the H3K27ac reads, whether or not they carried the somatic mutation genotype, as compared with the allele frequency of the same variant detected by WGS (P=6.2×10^−11^; **Figures 2D** and **S5D**). These results suggest that, in tumoral samples, differential histone modification enrichment at VREs is associated with the somatic mutation genotype. In the same line, and further confirming these results, we uncovered divergent H3K27ac enrichment at VREs and differential gene expression of nearby gene in samples mutated versus wild-type samples i.e., samples lacking somatic mutations in the VRE of interest (P=2.22×10^−16^) (**Figure S5E**). We observed examples of VREs associated with gains of H3K27ac, in mutated vs. non-mutated samples, proximal to induced genes implicated in the insulin secretion pathway (*PCSK1*^40^, *G6PC2*, *SLC30A8*^41^). On the other hand, VREs associated with reduced H3K27ac enrichments were proximal to downregulated genes implicated in p53 mediated apoptosis (e.g. *AEN*^42^, *DAB2IP*^43^ or *PHLDA3*^44^) and in critical components of the polycomb group complex (*RING1* and *JARID2*), whose function is that of maintaining a transcriptionally repressive chromatin state at specific genomic loci (**Figure 2E**).

Finally we uncovered that a significant fraction of somatic SNVs within VREs (37%) may affect the regulatory grammar by disrupting (eg. TP53) or creating (eg. SOX17) new TF binding motif sequences, thus providing a potential mechanism linking noncoding mutations to promotion of tumorigenesis (**Figure 2F**).

To provide a comprehensive overview of the mutational landscape in insulinomas, we combined all identified genetic alterations, both coding and noncoding, providing an extensive set of genes potentially implicated with the tumor development. (**Figure 2A** and **Table S4**; see Methods). This broad view allowed uncovering an expected enrichment of genes involved in cell cycle, cell growth and nervous development. Interestingly, genes encoding histone modifier enzymes were also found to be enriched within those mutated in insulinomas (**Figures S5F**). Overall, 92.5% of tumor samples bore a mutation affecting at least one gene listed as a *histone modifier* GO:0016570 (**Figures 2A**).

### Uncovering tumor-specific regulatory domains

Motivated by the observation of an enrichment of histone modifier enzymes within the genes mutated in insulinomas (**Figure 2A**), we sought to explore the genomic distribution of histone post-transcriptonal modifications in these tumors. Earlier studies demonstrated that large domains of H3K27ac underlie clusters of enhancers responsible for regulating key cell identity genes, having a functional role in disease susceptibility and cancer functions^22,45,46^. Furthermore, our interest was piqued by the correlation between the number of insulinoma-selective REs and gene up-regulation (**Figure 1E**), prompting us to explore the distribution of the newly mapped H3K27ac profiles along the genome. We found that insulinoma-selective H3K27ac sites were not evenly distributed throughout the genome (**Figure 3A**), but instead formed 375 clusters^22^ (**Figure S6A** and **Tables S3** and **S6**; see Methods), which we called **Insulinoma Regulatory Domains (IRD)**. These domains contained ∼50% of all H3K27ac sites gained in insulinomas and mirrored *super-enhancer* chromatin features, such as high enrichment in H3K27ac signal (**Figure S6B**) and stronger changes upon insulinoma transformation compared to other insulinoma-specific orphan regions (**Figure 3B**). Moreover, IRDs map in proximity of insulinoma-selective transcripts annotated to functions involving growth and transforming growth factor β (TGF-β) binding (**Figure 3C**). Other enriched terms were related to guanyl (ribo)nucleotide binding, mainly driven by the upregulation of genes from the GIMAP family, which are located within an IRD homogeneously present in the different insulinoma samples, and absent from control human islets (**Figure 3D**). Of note, overexpression of GIMAP genes has been implicated in T-cell leukemogenesis^47^. These data suggest that a subset of active enhancers are linked with tumor growth, opening the possibility to uncover driver regulatory mechanisms.

**Figure 3:**
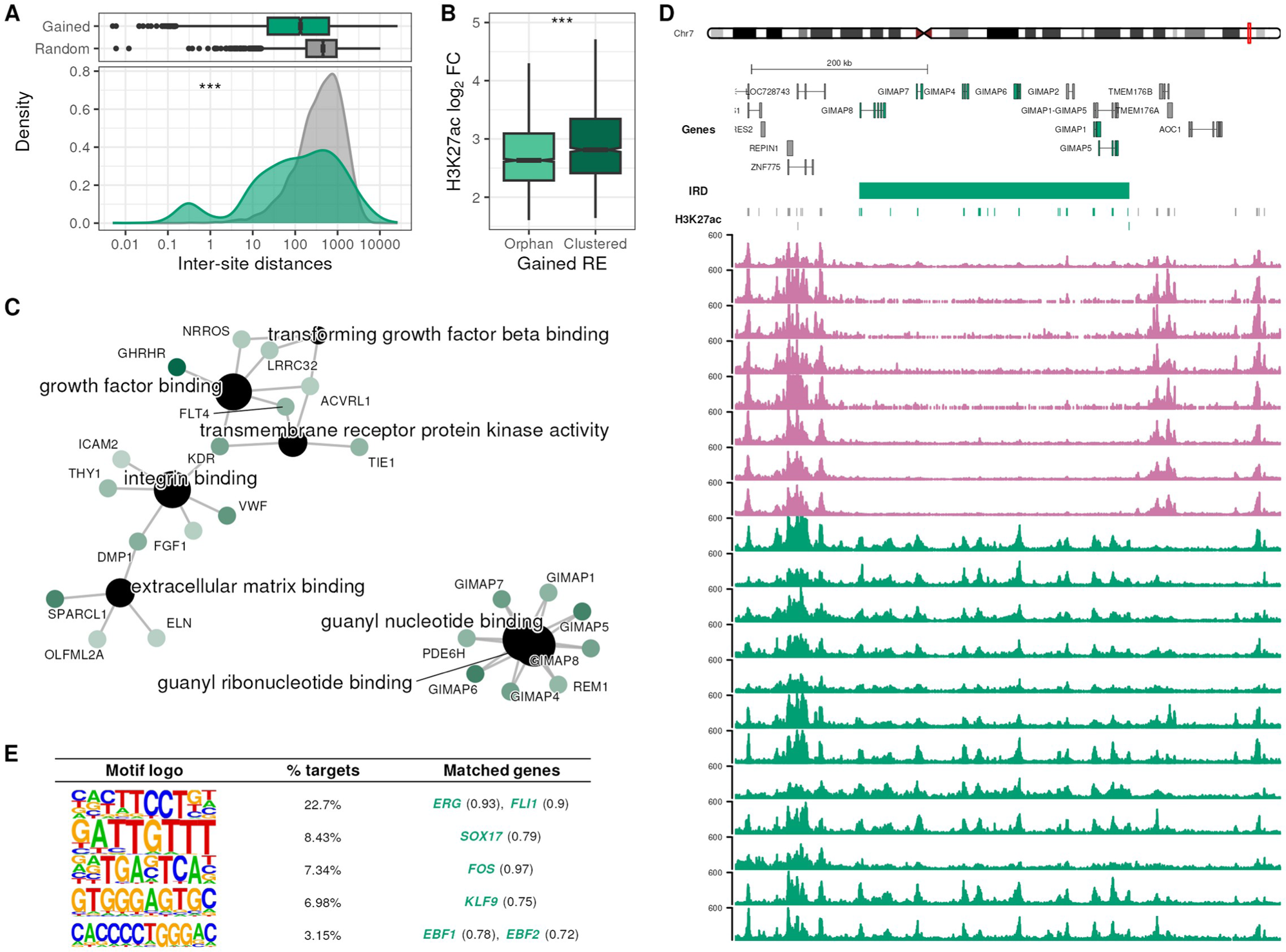
Insulinoma Regulatory Domains (IRD) consist of clusters of H3K27ac-selective sites. **A**, H3K27ac insulinoma-selective (gained) sites are highly clustered as their inter-site genomic distance is smaller than expected (random distribution, gray). Based on this analysis we defined 375 Insulinoma Regulatory Domains (IRD) (see Methods). **B**, Upon transition from normal β cell to insulinoma, REs located in IRD exhibit higher gains of H3K27ac enrichment than orphan RE. Two-sided Wilcoxon test ***P < 0.001. **C**, Gene Onthology: Molecular Function (GO:MF) annotation of up-regulated genes associated to IRD are related to growth factor binding, TGF-β pathway and guanyl nucleotide/ribonucleotide binding. Shade of green is proportional to the gene log2 fold change. **D**, Representative view of the *GIMAP* locus, encoding GTPases of the immunity-associated protein family (IMAP)^59^. *GIMAP* transcripts are induced in insulinomas and are encompassed by a large IRD composed of more than 15 sites consistently enriched of H3K27ac in 12 different insulinoma samples (green), but depleted of the active histone mark in untransformed human pancreatic islets (pink). **E**, Top *de novo* motifs identified by HOMER in nucleosome-free regions (NFR) at gained RE in IRDs. Only motifs present in more than 1% NFRs and matched to an up-regulated gene in insulinomas (score > 0.7) are shown.

Enhancer sequence stores information for the TFs potentially binding at accessible chromatin. To infer which TFs could be acting through IRDs and orchestrating neoplastic transition, we integrated open chromatin profiles from human pancreatic islets^23^ and 122 ENCODE cancer cell lines with H3K27ac sites at IRDs to identify putative nucleosome-free regions (NFR) (**Figure S6C**; see Methods). A *de novo* motif analysis identified 5 overrepresented motifs that matched TFs upregulated in insulinomas (**Figure 3D**), which may be driving transcriptional activity at IRDs. Such TFs include the ETS family (ERG/FLI1), SOX17, FOS, KLF9 and EBF1/2. Interestingly, many of these TFs not only seem to bind and regulate IRDs, but their own genes are also regulated by an IRD (**Figure S6D**). This observation matches the definition of Core Transcriptional Regulatory Circuitries (CRCs)^48^, in which TFs key for cell identity are inter-connected in regulatory loops through the super-enhancers that regulate their own expression. We thus sought to systematically infer sample-specific CRCs by producing patient-specific regulatory domains, uncovering that three motifs enriched in IRDs are part of CRCs in most insulinoma samples, namely *EBF1*, *ERG* and *SOX17* (**Figure S6E**). Of note, the SOX17 binding motif was also identified as being recurrently created by noncoding somatic variants in VREs (**Figure 2F**)

In summary, we observed insulinoma-specific activation of clustered RE (IRD) whose putative target genes are related to tumoral growth and harbor recurrent binding sites primarily for ETS and SOX17 TFs.

### Insulinoma Regulatory Domains map to polycomb repressed regions in healthy human islets

Motivated by two key observations: 1) the presence of a higher frequency of mutations affecting genes with histone modifier functions, and 2) the activation of extensive clusters of REs (IRDs) in insulinomas, we searched for potential chromatin-driven events in islet-cell tumor transition. To this end, we mapped IRDs to chromHMM chromatin states in both untransformed human pancreatic islets^49^ and a human β-cell line^50^. Most IRDs (40-80%) lie in regions annotated as quiescent or actively repressed by polycomb in the untransformed tissues (**Figure 4A** and **S7A**). To assess whether this overlap is statistically significant, we compared our data with newly generated and public H3K27me3 control tissue datasets^51–53^, a histone modification typically associated with polycomb repression. Indeed, we observed that IRDs displayed positive and significant z-scores (overlap permutation tests, n=500, P < 0.05) (**Figure 4B**), suggesting that these chromatin domains were repressed in β cells before undergoing tumor transformation. We named this subset of IRD-localized REs that are H3K27me3 repressed in control tissues **Derepressed Insulinoma Regulatory Domains** (**DeIRD**; **Table S3**). Of note, these observations are further supported by previous findings suggesting that β cell repressed genes are transcribed in insulinoma samples^3^.

**Figure 4:**
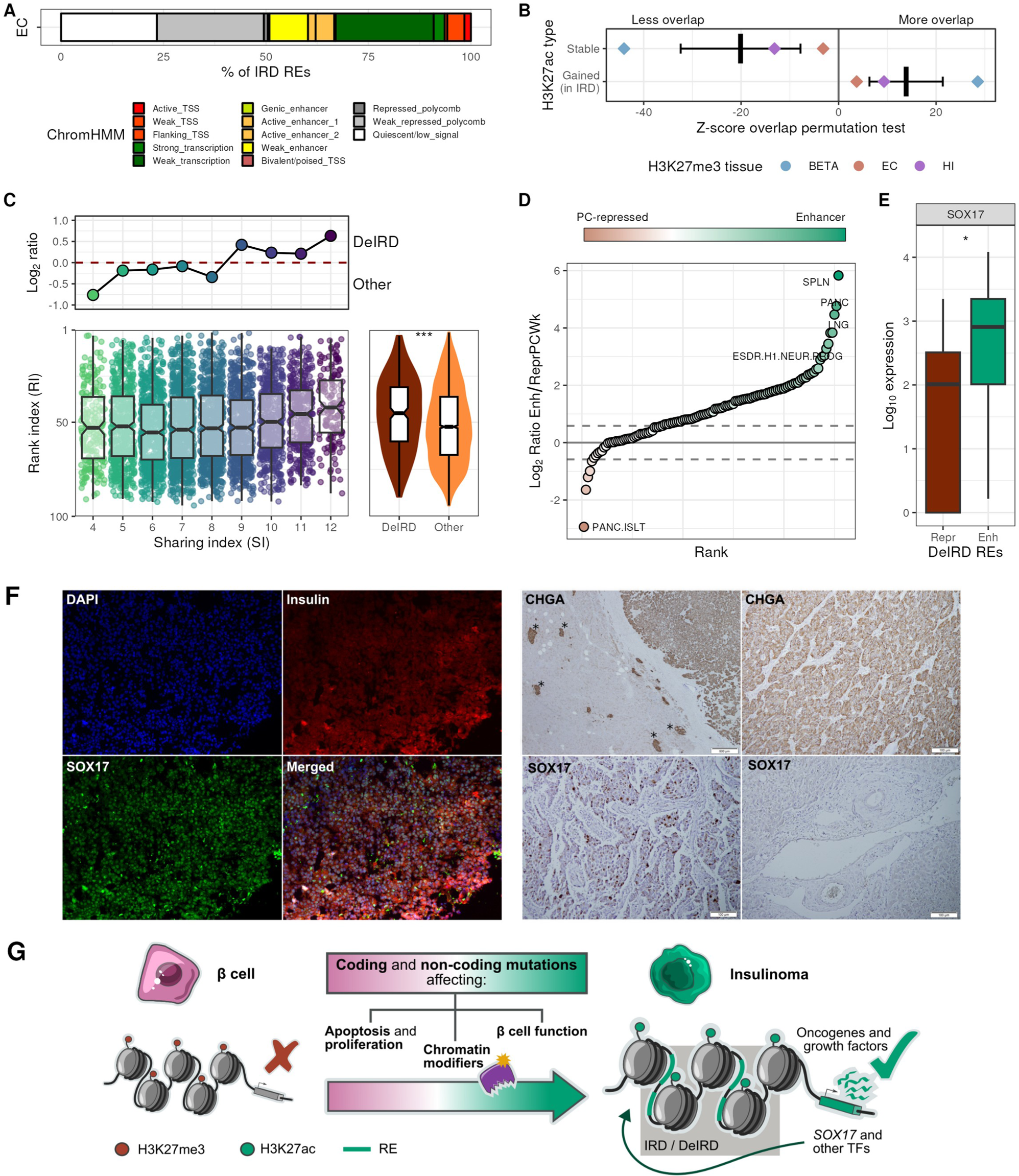
IRDs are polycomb-repressed in untransformed cell types. **A**, Chromatin state annotation of IRDs based on ChromHMM computed in EndoC-βH1. **B**, Distribution of permutation test z-scores comparing the overlap of IRD and stable REs in insulinomas with H3K27me3 peaks in β cells (BETA), EndoC-βH1 cells (EC) and human pancreatic islets (HI). Significant z-score are represented as diamonds, non-significant as dots. A black line depicts the mean z-score. **C**, Distribution of rank and sharing indexes in insulinoma-selective REs (gained). The top plot shows the ratio between DeIRDs and other gained REs at each sharing index value. The right plot shows the distribution of rank indexes of DeIRDs compared to thet of other gained REs. Two-sided Wilcoxon test: *** P < 0.001. **D**, Ratio between the number of DeIRDs annotated as enhancers (*Enh*) and as weak polycomb-repressed (ReprPCWk) in different tissues and cell lines from Epigenome Roadmap. **E**, Expression of *SOX17* in Epigenome Roadmap tissues in which DeIRDs are preferentially annotated as “polycomb-repressed” (Repr, ratio < −0.58) or as “enhancers” (Enh, ratio > 0.58). Two-sided Wilcoxon test: * P < 0.05. **F**, Immunofluorescence of an insulinoma sample showing co-staining of insulin and SOX17 (left). Insulinoma DAB immunohistochemistry staining (right) against Chromogranin A (CHGA) or SOX17. As a comparison, we show SOX17 in exocrine pancreatic tissue (bottom-right). Asterisks (*) mark healthy HI outside of the insulinoma. **G**, Simplified model illustrating the molecular mechanisms underlying insulinoma neoplastic transformation. Coding and noncoding mutations alter gene expression of chromatin modifiers resulting in the formation of IRDs and activation of polycomb-repressed regions. IRDs, control the expression of genes linked to neoplastic transformation and and other key TFs, such as SOX17. This reinforces a regulatory loop, thus facilitating the expression of genes that promote tumor cell growth and survival.

To measure whether DeIRDs could represent a common mechanism in the acquisition of an insulinoma phenotype, we took advantage of the sharing and rank indexes we computed for each H3K27ac site (**Figures 1F** and **S3C**). Interestingly we observed that DeIRDs were more clonal and more shared among patients, as compared to non-DeIRDs insulinoma REs (**Figure 4C**). These findings suggest that DeIRDs may represent a key mechanism to activate pathways implicated in the neoplastic transformation in insulinomas.

As DeIRDs are actively repressed in β cells, we next wondered whether these regions have enhancer functions in other tissues or cell types. To answer this question, we obtained all ChromHMM annotations available for all cell types and tissues in the Epigenome Roadmap^49^ and computed the ratio of DeIRDs overlapping enhancers vs polycomb-repressed regions. We confirmed that DeIRDs are preferentially annotated as polycomb-repressed in pancreatic islets (**Figure 4D**). Conversely, these same regions are annotated as enhancers in various cell types, including neural progenitors. This suggests that the pancreatic endocrine tumoral cells may exploit regulatory networks active in other cell populations. We wondered whether the activation of these regions could be orchestrated by the TFs identified to be acting at IRDs (**Figure 3E**). We therefore assessed their expression in all cell types annotated in the Epigenome Roadmap, categorized based on the DeIRD’s preferential annotation (polycomb-repressed or enhancer) in that specific cell type (**Figures 4E** and **S7B**). Interestingly, we observed that *SOX17, a* key developmental regulator, is induced in those tissues where DeIRDs are annotated as active enhancers. This observation suggests a pivotal role for this TF in driving insulinomas regulatory functions. Immunofluorescence and immunohistochemistry stainings confirm the nuclear localization of SOX17 at insulin-producing tumoral cells, excluding that the signal originates from other cell types such as endothelial cells or unaffected exocrine pancreas (**Fig. 4F**).

In summary, we propose a model in which coding and noncoding mutations alter chromatin modifiers, which in turn lead to activation of polycomb-repressed regions and the formation of IRDs. IRDs regulate the expression of genes related to neoplastic transformation (**Figure S7C-E**) and other key TFs, such as *SOX17* (**Figure S7F**), which reinforces a regulatory loop thus facilitating the expression of genes that promote tumor cell growth and survival (**Figure 4G**).

## Discussion

Characterization of the chromatin landscape of healthy human pancreatic islets has significantly contributed to shed light on the molecular mechanisms that underlie glucose metabolism-related diseases^54,55^. Much less is known on changes in regulatory function affecting human islets in disease state. In this study we charted a first draft of clinically relevant active regulatory regions shared between a large cohort of patient samples. Moreover, we characterized for the first time noncoding regulatory functions in insulin-producing neuroendocrine tumors.

We analyzed WGS somatic mutations derived from a large cohort of insulinomas. With the notable exception of *YY1*, we confirmed that recurrent coding mutations are rare. In line with results obtained in other cancer types^25,56^, we found that a large fraction of somatic mutations fall in the noncoding genome. Interpretation of these variants is difficult due to our limited knowledge of the regulatory code and the lack of understanding of disease-state noncoding functions. Furthermore, only a small fraction of these variants are expected to have a functional role in the acquisition of a neoplastic phenotype.

By integrating tumor-specific regulatory maps and insulinoma somatic mutations we observed that SNVs map preferentially to noncoding genomic sites active in insulinoma, correlating with the local H3K27ac signal and linked to genes primarily affecting the insulin secretion pathway as well as oncogenes and tumor suppressors.

Our data suggests a complex interplay between genetic mutations and epigenetic changes leading to a uniform tumoral phenotype. We found that coding and noncoding somatic mutations affect chromatin modifier genes, including histone demethylases and acetylases, as well as components of the polycomb and trithorax complexes. These alterations are coupled with extensive chromatin remodeling, which included the activation of large chromatin domains that are polycomb repressed in untransformed pancreatic islets. This pervasive reshape of noncoding regulatory functions leads to the activation of growth factors and oncogenes possibly driving the proliferative phenotype of the tumoral cells.

Our analyses identify key transcription factors playing a major role in orchestrating the regulatory changes leading to the insulinoma phenotype. Genetic and epigenetic data converge on the identification of *SOX17* having a driving role in the insulinoma phenotype: 1) the gene, which is repressed in normal human pancreatic islets, is up-regulated in insulinomas and its locus contains DeIRDs (**Figure S7F**); 2) its binding motif is recurrently created through somatic mutations (**Figure 2F**); 3) SOX17 binding sites are enriched throughout IRDs (**Figure 3E**); 4) tissues in which DeIRDs harbor active enhancers exhibit increased *SOX17* expression (**Figure 4E**). SOX17 is a key regulator of endoderm development which was shown to interact with the canonical Wnt signaling pathway^57^. In line with our findings, deletion of the polycomb-group protein EZH2 in human embryonic stem cells was shown to lead to derepression of developmental regulators including SOX17, resulting in self-renewal defects and misactivation of endoderm regulatory programs^58^.

Collectively our work implicates noncoding regulatory functions to the development of islet-cell derived tumors. By incorporating novel noncoding regulatory maps that encompass sequences critical to the loss of β cell identity and impaired insulin secretion, we can gain valuable insights into the essential functions involved. Further work will be needed to functionally delve into the impact of noncoding mutations in insulinomas and dissect those driving the neoplastic proliferation of human β cells. Newly defined regulatory maps and insulinoma somatic mutations can be visualized online along with other islet regulatory annotations (-*upon manuscript publication*-) at www.isletregulome.org.

## Acknowledgments

We thank Dr. Claudia Arnedo (IRB) for helpful discussions regarding the somatic mutation distribution across the non-coding genome. We acknowledge the patients and the Parc de Salut Mar MARBiobanc (PT20/00023) integrated in the Spanish National Biobanks Network from ISCIII for their collaboration.

## Author contributions

R.N., V.S., A.F.R., C.B.B., B.P.G., H.R. and S.P. performed wet-lab experiments. M.Ramos, M.S., R.N and G.F.P.. developed and performed bioinformatic analyses. M.F., L.Piemonti, R.C., E.M., M.N., S.P. supplied tissues. L.Pasquali, M.C., N.L., R.L., M.Rovira A.G. participated in the interpretation of the data and supervised the work. L.Pasquali designed the study. All authors read and approved the manuscript.

## STAR Methods

### RESOURCE AVAILABILITY

#### Lead contact

Further information and requests for resources and reagents should be directed to and will be fulfilled by the lead contact, Dr. Lorenzo Pasquali.

#### Materials availability

This study did not generate new unique reagents.

#### Data and code availability

RNA-seq, H3K27ac ChIP-seq and WGS data have been deposited at EGA and are publicly available as of the date of publication. Accession numbers are will be listed in the key resources table.

All original code has been deposited at Zenodo and will be publicly available as of the date of publication. DOIs will be listed in the key resources table.

Any additional information required to reanalyze the data reported in this paper is available from the lead contact upon request.

### EXPERIMENTAL MODEL AND STUDY PARTICIPANT DETAILS

Insulinoma samples were obtained from subjects who provided informed consent deposited at the Istituto Scientifico Ospedale San Raffaele, Milan, Italy. The study was approved under the protocol DiabPanc (P242/ER/mm) by the Ethical Committee of the Istituto Scientifico Ospedale San Raffaele. All experiments, conducted in accordance with the Declaration of Helsinki, were performed following procedures approved by the institutional research committees of the Institute for Health Science Research Germans Trias i Pujol, Barcelona, Spain (PI/15/051). All patients or their parents gave informed consent and all samples and data were handled protecting patients’ privacy. Patients sample details are provided in **Table S1.**

The samples were obtained upon removal of the tumour without interfering with the clinical patient management. A tumor mass measuring 4-6 cm^3^ was extracted from the resected insulinoma. A portion of each tumor mass was frozen for transcriptomic and genome sequence analysis, while another fraction was preserved using 1% formaldehyde to maintain the DNA-protein contacts for subsequent ChIP-seq assays. Whole blood was also collected from each patient, serving as a source of germline DNA. DNA was extracted using QIAamp DNA mini kit (Qiagen) according to the manufacturer’s instructions. DNA quality was assessed using Nanodrop (Thermo Fisher Scientific) and quantified using Qubit (Thermo Fisher Scientific) technology.

Additionally, 3 insulinoma tumor specimens from formalin-fixed paraffin-embedded blocks, were obtained from Parc de Salut MAR Biobank (MARBiobanc), Barcelona, Spain (2023S017E) and used for staining analyses.

Human pancreatic islet cells were isolated from donors in asystole or cardiorespiratory arrest (controlled asystole donation), without a history of alteration of the glucose metabolism, in accordance with national laws and institutional ethical requirements at the Hospital Universitari de Bellvitge, Barcelona, Spain. The cells were shipped in culture medium and then cultured for 72h before undergoing experimental procedures.

### METHOD DETAILS

#### RNA-seq

RNA was extracted from 11 frozen tumor samples using the AllPrep DNA/RNA extraction kit (Qiagen) resulting in >60 ng/μl yields per sample and RNA integrity numbers (RIN) >7.0. RNA libraries were prepared by ribosomal RNA depletion and were sequenced on a HiSeq 2000 platform (Illumina) to produce 150 bp paired-end reads with an average of 95 million reads per sample.

#### ChIP-seq and Cut&Tag

ChIP-seq was conducted using tagmentation (ChIPmentation), following a previously described method^60^. The samples fixed with 1% formaldehyde were sonicated to achieve an average fragment size of approximately 200bp. Immunoprecipitation was performed on 35μg of chromatin in a 0.4% SDS IP buffer using 1.5μl of anti-H3K27ac antibody (abcam ab4729) and 50μl of 10% BSA. After incubation, the immunoprecipitated DNA fragments were hybridized to protein A+G beads and then washed using low-salt, high-salt, and LiCl wash buffers. Subsequently, the IPs underwent a 10-minute incubation with 1μl of Tagment DNA enzyme, followed by additional washes with RIPA and Tris-EDTA buffers. To generate ChIP libraries, elution was performed using a 1% SDS, 0.1M NaHCO3 buffer. Two μl of each library were amplified in a 10 μl qPCR reaction containing 0.15 μM primers, 1 × SYBR Green and 5 μl NEBNext High-Fidelity 2X PCR Master Mix (NEB M0541S), to estimate the optimum number of enrichment cycles needed for library amplification. The libraries were then amplified using the Nextera DNA Library Prep Kit (15028212, Illumina, San Diego, USA). Semi-quantitative PCR assays at target positive and negative control sites were performed to estimate the efficiency of the ChIP experiment before sequencing (data not shown). Finally, sequencing of the ChIP libraries was carried out using a single-end protocol with 50bp reads, with a minimum of 30 million.

Cut&Tag was used to profile H3k27me3 in HI and EC and was performed as previously described^61^. For human islets, we first disaggregated them into single cells using trypsin. All incubations at 4°C or room temperature were performed on a rotating wheel. Briefly, cells were harvested, counted, and washed in Wash Buffer. During washes, we activated the ConA coated magnetic beads by washing them with Binding buffer and resuspended in 1 volume of binding buffer before incubation with the cells. In each experiment, 100,000 cells and 10 μL of beads were used. Next, bead-bound cells were resuspended in 50 μL ice-cold Antibody buffer and transferred to a LoBind tube. Primary antibody against H3K27me3 (Millipore, #07-449) was added 1:100 and incubated overnight at 4°C. Afterwards, samples were incubated with the secondary antibody (Antibodies Online, #ABIN101961) diluted 1:100 in Dig-wash buffer and incubated at room temperature for 1h. Cells were washed with Dig-wash buffer and incubated with a mix of pA-Tn5 adapter complex (Cutana, #15-1017) in Dig-300 buffer at room temperature for 1h. After incubation, beads were washed with Dig-300 buffer, resuspended in 300 μL of Tagmentation buffer and incubated at 37°C for 1h. Then, the tagmentation was stopped and decrosslinking was performed before DNA purification. The DNA was purified by phenol-chloroform extraction and 21 μL were used for library amplification, performed as described in the original protocol. Post-PCR clean-up was performed by adding 1.3Xof Ampure XP beads (Agencourt AMPureXP, Beckman-Coulter, #A63880) and samples were eluted in 25 μL 10 mM Tris-HCl pH 8.

#### Immunofluorescence and immunohistochemistry stainings

Insulinoma samples were embedded in OCT. Samples were sectioned at 4-5 uM prior to immunostaining. For Immunofluorescence stainings, tissue sections were fixed with 4% PFA for 20min at 4°C, washed with PBS and blocked in 0.5% (v/v) triton PBS, 1% (v/v) FBS PBS for 30min at room temperature. Incubation with primary antibodies (anti-Insulin: DAKO A0564, anti-FLI1: Thermo PA5-29597, anti-SOX17: Neuromics GT15094, anti-FGF1: Abcam ab9588) was performed overnight at 4°C in 0.5% (v/v) triton PBS. Samples were washed with PBS for 30min at 4°C before incubation with secondary antibodies for 2h at 4°C. Lastly, samples were stained for nuclei with DAPI.

For immunohistochemical stainings, tissue sections were dewaxed and rehydrated (2x xylol 5min, 2x EtOH 100% 5min, 2x EtOH 96% 3min, EtOH 70% 3min, 5min dH2O) before performing antigel retrieval in ph6 citrate buffer in a decloaking chamber. Then, tissue sections were permeabilized with 0.25% (v/v) triton PBS for 20min at room temperature and blocked in 1% (v/v) donkey-serum for 30min at room temperature. Incubation with primary antibodies was performed overnight at 4°C prior to incubation with secondary-HRP conjugated antibody for 1h at RT. Staining was performed with DAB Peroxidase Substrate Kit (Liquid DAB+Substrate Chromogen System, Dako), following manufacturers instruction.

Epifluorescent images were acquired on a ZeissAxio_ObserverZ1_Apotome inverted fluorescent microscope. Brightfield images were taken on a LeicaZ16 APO stereomicroscope.

### QUANTIFICATION AND STATISTICAL ANALYSIS

#### RNA-seq

Reads were aligned to gencode version 38 using Salmon^62^ (version 1.3.0). We loaded the results into R using tximport^63^ (version 1.26.0), summarized transcript information into genes and kept protein-coding genes for downstream analyses. We also downloaded published RNA-seq samples from insulinomas and from human islets and beta cells to use as comparison, which were processed in the same manner (**Table S7**).

#### ChIP-seq and Cut&Tag

Reads were mapped to hg38 using Bowtie2^64^ (version 2.4.1) with the “--local” parameter in the case of ChIP-seq and with “--very-sensitive --no-mixed --no-discordant --phred33 -I 10 -x 100”, in the case of Cut&Tag. Next, we removed duplicates and reads mapping to non-cannonical chromosomes or to ENCODE blacklisted regions using Samtools^65^ (version 1.10). We performed peak calling using MACS2 (version 2.2.7.1) with arguments “--broad --broadcutoff 0.1 --nomodel”.

Additionally to the data generated in the current study, publicly available H3K27ac ChIP-seq raw data from human islets, other cancers, cell lines and normal tissues were downloaded and processed in the same manner. The full list with references of the employed datasets are listed in **Table S7**.

#### Consensus H3K27ac sites

We retained all peaks called with -log_10_ p-value > 4 and used the R package DiffBind^66^ (version 3.8.3) to create **consensus peaksets** in different ways:

1. **Stringent consensus dataset**: The consensus peaksets for the comparison of insulinomas vs human islets, and the diverse H3K27ac heatmaps were obtained by selecting in a tissue-specific manner those regions present in more than 30% of the samples. Then, tissue-specific consensus peaks were merged together.
2. **Comprehensive consensus dataset**: The consensus peakset used for the variance analysis (**Figure S3B**), as well as the region set that was contrasted with the insulinoma somatic SNVs in order to identify VREs, were created by merging together all peaks in all samples, without filtering for recurrence.

To obtain the number of peaks in each peak we employed the function *featureCounts()* from the Rsubread^67^ R package (version 2.12.3).

#### Differential analysis of RNA-seq and ChIP-seq

To perform differential analyses in our genomic data we used DESeq2 ^68^ with default parameters (version 1.38.1). Genes/regions were considered significantly **gained** when adjusted p-value < 0.05 and log2FC > 1, significantly **lost** when adjusted p-value < 0.05 and log2FC < −1 and **stable** if they did not pass these thresholds. Of note, all downstream analyses were performed on insulinoma-selective genes and sites (gained), thus excluding signals that could arise from non β-cell types populating the normal endocrine pancreas.

#### Data normalization and transformation

When needed, sequencing data was transformed using the variance stabilizing transformation (VST) method implemented as the vst() function in the DESeq2^68^ R package with default parameters.

#### Assigning regulatory elements to target genes

##### Chromatin analyses

- **Figure 1E**: To assign RE to putative target genes in an unbiased manner, we designed a hybrid approach that used fixed-size windows (40kb around TSS) and double elite interactions present in the GeneHancer database^69^.
- **Figure 3C**: REs were annotated using fixed-size windows (40kb around TSS) to genes up-regulated in insulinomas compared to human islets.

###### VREs

All VREs were annotated to genes whose TSS was closer than 5 kb upstream and 1 kb downstream. Additionally, they were annotated to the nearest upstream and downstream gene TSS within a 1 Mb distance (similar to the default algorithm used in GREAT^70^).

#### Sharing and Rank Indexes

Sharing and rank indexes were obtained as previously described^26^. The code is implemented in the function *get_ranking_sharing_index()* in our custom meowmics R package (https://github.com/mireia-bioinfo/meowmics).

#### Whole-genome sequencing

Illumina NovaSeq 6000 technology was used to sequence whole-genome 150 bp paired-end TruSeq PCR-free libraries. The raw sequencing data was aligned with BWA-MEM (version 0.7.17) to the NCBI Human Reference Genome Build hg38. Duplicates were marked using samblaster (version 0.1.24) and BAMs were sorted and indexed using Samtools^65^ (version 1.9). Samtools depth was employed to calculate alignment and coverage metrics, revealing a mean read depth of 37x±5x for peripheral blood cells and 58x±7x for tumor samples.

Additionally to the 26 whole genome sequence (WGS) generated in this study, the raw reads of 14 WGS from Scarpa et al.^71^, 20 whole exome sequence (WES) from Wang et al.^72^ and 20 WES from Cao et al.^73^ including insulinoma and matched blood samples were downloaded and processed in the same way in order to enable an homogeneous variant calling over 40 insulinoma samples.

#### Somatic variant calling and filtering

SNVs (Single Nucleotide Variants) and INDELs (Insertions and Deletions) were identified using Strelka2^74^ (version 2.9.10) and Mutect2^75^ (version 2.23.0) with default settings. Tumor and matched blood sequencing data were used to remove germinal variants. Following the recommended best practice, Strelka2 was executed incorporating the candidate indel generated by manta^76^ (version 1.6.0). Only “PASS” variants in Variant Call Format (VCFs) resulting from both callers algorithms were included. Variants present in cohort-specific Panel of Normals and in gnomAD^77^ (version 3.1) with a VAF > 0.001 were removed. Additionally, variants within UCSC Common set. dbSNP150^78^ and/or segmental duplications, simple repeats and masked regions were also excluded. The resulting VCFs were annotated using ANNOVAR^79^ (version 2020Jun08) (**Figure S4A**).

We next used IntOGen^36^, to discover insulinoma coding driver genes by inferring signals of positive selection as previously described.

Insulinomas single nucleotide somatic mutations (n=25.497), were mapped to a consensus datasets of H3K27ac sites active in insulinomas and its normal tissue counterpart (see “ChIP-seq and Cut&Tag” section) resulting in 1,640 insulinoma variant regulatory elements (VREs).

#### Tumor mutational burden (TMB)

ANNOVAR annotated VCF were converted to MAF using the annovarToMaf from maftools^80^ R package (version 2.16.0). tcgaCompare maftools function was used to calculate the TMB of all the datasets. Comparative plots were generated using the tcgaCompare function.

#### Mutational signatures

Extraction of SBS and ID signatures was performed using SigProfilerExtractor^33^ (version 1.1.4), which employs NMF to extract optimal number of mutational signatures in a given cohort of tumors. Signatures were extracted the *novo* and decomposed based on Catalog of Somatic Mutations in Cancer (COSMIC)^81^ (version 3) using a cosine similarity greater than 0.9.

#### Structural variant calling

SVs were called using Delly^82^ (version 1.1.6), GRIDSS^83^ (version 2.11.1-1), Manta^76^ (version 1.6.0), Smoove (https://github.com/brentp/smoove) (version 0.2.8) in a somatic configuration. GRIDSS SVs were post-annotated as DEL, DUP, INV, INS and BND with sv_type_infer_gridss.R script provided in GRIDSS. “PASS” variants were annotated using Duphold^84^ based on Duphold’s flanking fold change (DHFFC) annotation. Deletions with a DHFFC value greater than 0.7 and duplications with a value less than 1.25 were excluded. ENCODE DAC blacklist was used to remove regions with anomalous, unstructured and high signal/read counts. “PASS” variants identified in all 4 callers were included.

Population SVs were inferred from 1,000G catalog. To this end 2,504 low-coverage BAMs were downloaded from the 1,000 genomes AWS S3 bucket (s3://1000genomes/phase3/data/) to build a control reference STIX database using excord (version 0.2.4) (https://github.com/brentp/excord), giggle^85^ (version 0.6.3) and STIX^85^ (version 1.0). First, SV alignment evidence was extracted from BAM using excord, considering both discordand and split reads (discordant distance=500). Next indexes for each excord evidence were generated by giggle. Finally, the database was created using STIX as described elsewhere^85^. The same procedure was applied to the patient sample cohort to obtain an insulinoma STIX database.

We contrasted “PASS” SV variants obtained from the two datasets and removed SVs with evidence of >10 counts in 1000 Genomes (considered population variants) as well as those with >1 counts in the insulinoma control cohort (considered germline SV).

Samplot^86^ (version 1.3.0) was used to generate the images for each SV and remaining SVs variants were manually curated creating the final SV dataset.

SVs were annotated using the annotation from GENCODE (release 18)^87^. SVs affecting coding sequences were annotated as CDS SVs. non-CDS SV were assigned to genes with a TSS at <1Mb distance.

#### Variants enrichment analysis

We used the regioneR^88^ R package v.1.32.0 to assess the enrichment of insulinoma single nucleotide somatic mutations in relation to overlapping H3K27ac enriched sites mapped in insulinomas, as well as in other cancer types, primary tissue types, or cell lines. The H3K27ac raw data from all datasets underwent uniform processing, as detailed in the “ChIP-seq” methods section.

A null distribution was generated by permuting 500 times a set of regions matched in size and structure to the original H3K27ac region dataset. The number of insulinoma single nucleotide somatic mutations overlapping the H3K27ac dataset was computed and compared with the intersections obtained with the matched null distribution. The enrichment score was determined by measuring the number of standard deviations that the overlapping count differs from the median of the null overlapping count. Subsequently, the exact p-value was computed by fitting a density function to the null distribution obtained from the matched random variant set. Finally, this p-value underwent correction for multiple testing using the Benjamini–Hochberg method. Enrichments or depletions with a Benjamini–Hochberg-adjusted P <0.05 were considered statistically significant and with marked as red dots (**Figure 2B**).

#### Differential H3K27ac enrichment at VREs and nearby gene expression analysis

In each sample the number of aligned reads containing either the REF or ALT alleles of a SNV was determined for each H3K27ac-WGS matched BAM pair using Rsamtools^89^ (version 2.16.0). Only SNV overlapped by a minimum of 4 H3K27ac reads were retained. Reads retrieved from WGS and H3K27ac were used to separately calculate the absolute log2FCs between reads of REF and ALT alleles (**Figure 2D**).

Differential H3K27ac enrichment was computed at each VRE computing the absolute log2FC H3K27ac signal between mutated vs. wildtype samples. As a control, the same calculation was performed after randomizing the sample genotypes (number of permutation=500). RNA differential analysis was conducted in a similar manner. In this case, RNA-seq reads from the closest VRE transcript were used to calculate absolute log2FC (**Figure S5E**).

#### Conservation analysis

Peaks were extended from the center 1Kb to each direction. Mean phylogenetic conservation scores were computed over 20 bp segments, using values obtained from the phastCons100way dataset^90^ (**Figures S2F** and **S5E**).

#### Oncoplot

Somatic variants, including SNVs/INDELs, VREs and SVs (described in “Somatic variant calling and filtering”, “Structural variant calling” and “VREs and gene annotation”) were summarized using custom scripts and plotted into an oncoplot using the ggplot2^91^ (version 3.4.3) R package. Genes with mutations observed in a minimum of three samples and/or exhibiting a recurrent VRE (same regulatory element affected across multiple samples) were included in the heatmap. Driver coding mutations, denoted with a dot, were inferred by IntOGen as described in “Somatic variant calling and filtering”.

#### Pathway enrichment analyses

GSEA of up-regulated genes in insulinomas was performed using the fgsea ^92^ R package (version 1.24.0) with MSigDB annotations (version 7.4).

Pathway enrichment analysis of regulatory elements was conducted using rGREAT^70^ (version 2.2.0) R package. The hallmark gene sets were retrieved from the Molecular Signatures Database (MSigDB) using msigdb:H rGREAT collection.

Enrichment of Gene Ontology Molecular Function (GO-MF) terms in gained genes associated to IRD and VREs was assessed using the function enrichGO from the clusterProfiler^93^ R package (version 4.6.2). The results were plotted using the cnetplot function from the enrichplot^94^ R package (version 1.18.4).

#### Insulinoma Regulatory Domains

Regulatory Domains were identified as previously described^22^. The code is implemented in the function *get_enhancer_clusters()* in our custom meowmics R package (https://github.com/mireia-bioinfo/meowmics). Briefly, we selected gained H3K27ac sites and randomized them within their chromosome to derive chromosome-specific thresholds (10th percentile of the random distribution) for stitching together H3K27ac sites into domains. We selected domains containing at least 3 H3K27ac sites.

#### Sequence composition and transcription factor analyses

A collection of ATAC-seq data from 122 human cell lines obtained from the ENCODE portal and including experiments performed on human islets cells from a previous publication^23^, were used to generate a comprehensive database of regions of open chromatin representative of different human cell types. Each peak summit was extended 200bp upstream and downstream from the center to define Nucleosome Free Regions (NFRs).

NFRs overlapping distal REs in IRDs (**Figure 3E**) were used as input for *de novo* motif analysis with HOMER^95^ (version 4.11) findMotifGenome.pl tool, using parameters ‘-size 200 -mask -preparsed’. Matched genes for the overrepresented sequences were selected as described in the figure legends. The same parameters were used to infer *de novo* TF binding motif in VREs (**Figure S5C).**

Prediction of TF binding sites disrupted or created by SNVs in VREs was performed using motifbreakR^96^ (version 2.14.2) R package and the HOMER motif data source from the MotifDb^97^ (version 1.42.0).

#### Core transcriptional regulatory circuitry

Interconnected circuitries of transcription factors acting in insulinomas were generated using the CRCmapper^48^. H3K27ac ChIP-seq reads, H3K27ac sites, and sample-specific IRDs were used as input. Briefly, the algorithm first assigns clusters of enhancers to the closest gene predicted to be expressed. Then, identifies the candidate core transcription factors from the genes linked to IRDs, and subsequently performs a known motif analysis using the H3K27ac sites. Finally, identifies auto-regulated TFs, and those that are binding IRDs generating a fully interconnected auto-regulatory loop. The original code has been edited to support genome GRCh38 build and alternative islet-specific TFs motif matrices were included in the database of positional weight matrices (PWMs). The improved code version, also implemented as a Singularity image, has been deposited at: https://github.com/mireia-bioinfo/CRCmapper.

#### Overlap with H3K27me3 regions

Overlap between individual H3K27me3 peaks (publicly available^51–53^ or generated in the present study, processed as described in “ChIP-seq and Cut&Tag”) and the RE composing the IRDs was computed. To contrast expected and observed overlap, we resampled the annotation coordinates 500 times using regioneR^88^ R package (version 1.30.0). We annotated as DeIRDs those RE that overlapped at least one H3K27me3 dataset.

#### Epigenome Roadmap ChromHMM and gene expression data

ChromHMM annotation and processed H3K27me3 peak files from HI^49^ and EC^50^ were downloaded from the respective sources. The function liftOver() from rtracklayer^98^ R package (version 1.58.0) was used to convert the coordinates to hg38. ChromHMM datasets were mapped to the RE composing IRDs.

15-state ChromHMM data from the Epigenome Roadmap^49^ were downloaded from their source (egg2.wustl.edu). We selected states “7_Enh” to represent “enhancers” and “14_ReprPCWk” to represent “polycomb-repressed” regions.

Gene expression from 57 epigenomes was downloaded from the same source. Counts were normalized using DESeq2^68^.

## References

1. Roy, N., and Hebrok, M. (2015). Regulation of Cellular Identity in Cancer. Dev. Cell 35, 674–684. 10.1016/j.devcel.2015.12.001.

2. Okabayashi, T. (2013). Diagnosis and management of insulinoma. World J. Gastroenterol. 19, 829. 10.3748/wjg.v19.i6.829.

3. Wang, H., Bender, A., Wang, P., Karakose, E., Inabnet, W.B., Libutti, S.K., Arnold, A., Lambertini, L., Stang, M., Chen, H., et al. (2017). Insights into beta cell regeneration for diabetes via integration of molecular landscapes in human insulinomas. Nat. Commun. 8, 767. 10.1038/s41467-017-00992-9.

4. Klöppel, G. (2011). Classification and pathology of gastroenteropancreatic neuroendocrine neoplasms. Endocr. Relat. Cancer 18, S1–S16. 10.1530/ERC-11-0013.

5. Cao, Y., Gao, Z., Li, L., Jiang, X., Shan, A., Cai, J., Peng, Y., Li, Y., Jiang, X., Huang, X., et al. (2013). Whole exome sequencing of insulinoma reveals recurrent T372R mutations in YY1. Nat. Commun. 4, 2810. 10.1038/ncomms3810.

6. Cromer, M.K., Choi, M., Nelson-Williams, C., Fonseca, A.L., Kunstman, J.W., Korah, R.M., Overton, J.D., Mane, S., Kenney, B., Malchoff, C.D., et al. (2015). Neomorphic effects of recurrent somatic mutations in *Yin Yang 1* in insulin-producing adenomas. Proc. Natl. Acad. Sci. 112, 4062–4067. 10.1073/pnas.1503696112.

7. Modali, S.D., Parekh, V.I., Kebebew, E., and Agarwal, S.K. (2015). Epigenetic Regulation of the lncRNA MEG3 and Its Target c-MET in Pancreatic Neuroendocrine Tumors. Mol. Endocrinol. 29, 224–237. 10.1210/me.2014-1304.

8. Karakose, E., Wang, H., Inabnet, W., Thakker, R.V., Libutti, S., Fernandez-Ranvier, G., Suh, H., Stevenson, M., Kinoshita, Y., Donovan, M., et al. (2020). Aberrant methylation underlies insulin gene expression in human insulinoma. Nat. Commun. 11, 5210. 10.1038/s41467-020-18839-1.

9. Flores, M.A., and Ovcharenko, I. (2018). Enhancer reprogramming in mammalian genomes. BMC Bioinformatics 19, 316. 10.1186/s12859-018-2343-7.

10. Huyghe, A., Trajkova, A., and Lavial, F. (2023). Cellular plasticity in reprogramming, rejuvenation and tumorigenesis: a pioneer TF perspective. Trends Cell Biol., S0962892423001575. 10.1016/j.tcb.2023.07.013.

11. The ICGC/TCGA Pan-Cancer Analysis of Whole Genomes Consortium, Aaltonen, L.A., Abascal, F., Abeshouse, A., Aburatani, H., Adams, D.J., Agrawal, N., Ahn, K.S., Ahn, S.-M., Aikata, H., et al. (2020). Pan-cancer analysis of whole genomes. Nature 578, 82–93. 10.1038/s41586-020-1969-6.

12. Comfort, N. (2015). Genetics: We are the 98%. Nature 520, 615–616. 10.1038/520615a.

13. Dietlein, F., Wang, A.B., Fagre, C., Tang, A., Besselink, N.J.M., Cuppen, E., Li, C., Sunyaev, S.R., Neal, J.T., and Van Allen, E.M. (2022). Genome-wide analysis of somatic noncoding mutation patterns in cancer. Science 376, eabg5601. 10.1126/science.abg5601.

14. Cnop, M., Abdulkarim, B., Bottu, G., Cunha, D.A., Igoillo-Esteve, M., Masini, M., Turatsinze, J.-V., Griebel, T., Villate, O., Santin, I., et al. (2014). RNA Sequencing Identifies Dysregulation of the Human Pancreatic Islet Transcriptome by the Saturated Fatty Acid Palmitate. Diabetes 63, 1978–1993. 10.2337/db13-1383.

15. Eizirik, D.L., Sammeth, M., Bouckenooghe, T., Bottu, G., Sisino, G., Igoillo-Esteve, M., Ortis, F., Santin, I., Colli, M.L., Barthson, J., et al. (2012). The Human Pancreatic Islet Transcriptome: Expression of Candidate Genes for Type 1 Diabetes and the Impact of Pro-Inflammatory Cytokines. PLOS Genet. 8, e1002552. 10.1371/journal.pgen.1002552.

16. Gonzalez-Duque, S., Azoury, M.E., Colli, M.L., Afonso, G., Turatsinze, J.-V., Nigi, L., Lalanne, A.I., Sebastiani, G., Carré, A., Pinto, S., et al. (2018). Conventional and Neo-antigenic Peptides Presented by β Cells Are Targeted by Circulating Naïve CD8+ T Cells in Type 1 Diabetic and Healthy Donors. Cell Metab. 28, 946–960.e6. 10.1016/j.cmet.2018.07.007.

17. Morán, I., Akerman, İ., van de Bunt, M., Xie, R., Benazra, M., Nammo, T., Arnes, L., Nakić, N., García-Hurtado, J., Rodríguez-Seguí, S., et al. (2012). Human β Cell Transcriptome Analysis Uncovers lncRNAs That Are Tissue-Specific, Dynamically Regulated, and Abnormally Expressed in Type 2 Diabetes. Cell Metab. 16, 435–448. 10.1016/j.cmet.2012.08.010.

18. Ackermann, A.M., Wang, Z., Schug, J., Naji, A., and Kaestner, K.H. (2016). Integration of ATAC-seq and RNA-seq identifies human alpha cell and beta cell signature genes. Mol. Metab. 5, 233–244. 10.1016/j.molmet.2016.01.002.

19. Arda, H.E., Li, L., Tsai, J., Torre, E.A., Rosli, Y., Peiris, H., Spitale, R.C., Dai, C., Gu, X., Qu, K., et al. (2016). Age-Dependent Pancreatic Gene Regulation Reveals Mechanisms Governing Human β Cell Function. Cell Metab. 23, 909–920. 10.1016/j.cmet.2016.04.002.

20. Blodgett, D.M., Nowosielska, A., Afik, S., Pechhold, S., Cura, A.J., Kennedy, N.J., Kim, S., Kucukural, A., Davis, R.J., Kent, S.C., et al. (2015). Novel Observations From Next-Generation RNA Sequencing of Highly Purified Human Adult and Fetal Islet Cell Subsets. Diabetes 64, 3172–3181. 10.2337/db15-0039.

21. Parker, S.C.J., Stitzel, M.L., Taylor, D.L., Orozco, J.M., Erdos, M.R., Akiyama, J.A., van Bueren, K.L., Chines, P.S., Narisu, N., NISC Comparative Sequencing Program, et al. (2013). Chromatin stretch enhancer states drive cell-specific gene regulation and harbor human disease risk variants. Proc. Natl. Acad. Sci. 110, 17921–17926. 10.1073/pnas.1317023110.

22. Pasquali, L., Gaulton, K.J., Rodríguez-Seguí, S.A., Mularoni, L., Miguel-Escalada, I., Akerman, İ., Tena, J.J., Morán, I., Gómez-Marín, C., van de Bunt, M., et al. (2014). Pancreatic islet enhancer clusters enriched in type 2 diabetes risk-associated variants. Nat. Genet. 46, 136–143. 10.1038/ng.2870.

23. Ramos-Rodríguez, M., Raurell-Vila, H., Colli, M.L., Alvelos, M.I., Subirana-Granés, M., Juan-Mateu, J., Norris, R., Turatsinze, J.-V., Nakayasu, E.S., Webb-Robertson, B.-J.M., et al. (2019). The impact of proinflammatory cytokines on the β-cell regulatory landscape provides insights into the genetics of type 1 diabetes. Nat. Genet. 51, 1588–1595. 10.1038/s41588-019-0524-6.

24. Cejas, P., Drier, Y., Dreijerink, K.M.A., Brosens, L.A.A., Deshpande, V., Epstein, C.B., Conemans, E.B., Morsink, F.H.M., Graham, M.K., Valk, G.D., et al. (2019). Enhancer signatures stratify and predict outcomes of non-functional pancreatic neuroendocrine tumors. Nat. Med. 25, 1260–1265. 10.1038/s41591-019-0493-4.

25. Corona, R.I., Seo, J.-H., Lin, X., Hazelett, D.J., Reddy, J., Fonseca, M.A.S., Abassi, F., Lin, Y.G., Mhawech-Fauceglia, P.Y., Shah, S.P., et al. (2020). Non-coding somatic mutations converge on the PAX8 pathway in ovarian cancer. Nat. Commun. 11, 2020. 10.1038/s41467-020-15951-0.

26. Patten, D.K., Corleone, G., Győrffy, B., Perone, Y., Slaven, N., Barozzi, I., Erdős, E., Saiakhova, A., Goddard, K., Vingiani, A., et al. (2018). Enhancer mapping uncovers phenotypic heterogeneity and evolution in patients with luminal breast cancer. Nat. Med. 24, 1469–1480. 10.1038/s41591-018-0091-x.

27. Ye, B., Fan, D., Xiong, W., Li, M., Yuan, J., Jiang, Q., Zhao, Y., Lin, J., Liu, J., Lv, Y., et al. (2021). Oncogenic enhancers drive esophageal squamous cell carcinogenesis and metastasis. Nat. Commun. 12, 4457. 10.1038/s41467-021-24813-2.

28. Li, Q.-L., Lin, X., Yu, Y.-L., Chen, L., Hu, Q.-X., Chen, M., Cao, N., Zhao, C., Wang, C.-Y., Huang, C.-W., et al. (2021). Genome-wide profiling in colorectal cancer identifies PHF19 and TBC1D16 as oncogenic super enhancers. Nat. Commun. 12, 6407. 10.1038/s41467-021-26600-5.

29. Stelloo, S., Nevedomskaya, E., Kim, Y., Schuurman, K., Valle-Encinas, E., Lobo, J., Krijgsman, O., Peeper, D.S., Chang, S.L., Feng, F.Y.-C., et al. (2018). Integrative epigenetic taxonomy of primary prostate cancer. Nat. Commun. 9, 4900. 10.1038/s41467-018-07270-2.

30. Australian Pancreatic Cancer Genome Initiative, Scarpa, A., Chang, D.K., Nones, K., Corbo, V., Patch, A.-M., Bailey, P., Lawlor, R.T., Johns, A.L., Miller, D.K., et al. (2017). Whole-genome landscape of pancreatic neuroendocrine tumours. Nature 543, 65–71. 10.1038/nature21063.

31. Jameson, J.L. ed. (2016). Endocrinology: adult & pediatric 7th edition. (Elsevier Saunders).

32. Ellrott, K., Bailey, M.H., Saksena, G., Covington, K.R., Kandoth, C., Stewart, C., Hess, J., Ma, S., Chiotti, K.E., McLellan, M., et al. (2018). Scalable Open Science Approach for Mutation Calling of Tumor Exomes Using Multiple Genomic Pipelines. Cell Syst. 6, 271–281.e7. 10.1016/j.cels.2018.03.002.

33. Bergstrom, E.N., Huang, M.N., Mahto, U., Barnes, M., Stratton, M.R., Rozen, S.G., and Alexandrov, L.B. (2019). SigProfilerMatrixGenerator: a tool for visualizing and exploring patterns of small mutational events. BMC Genomics 20, 685. 10.1186/s12864-019-6041-2.

34. Tate, J.G., Bamford, S., Jubb, H.C., Sondka, Z., Beare, D.M., Bindal, N., Boutselakis, H., Cole, C.G., Creatore, C., Dawson, E., et al. (2019). COSMIC: the Catalogue Of Somatic Mutations In Cancer. Nucleic Acids Res. 47, D941–D947. 10.1093/nar/gky1015.

35. Driehuis, E., Van Hoeck, A., Moore, K., Kolders, S., Francies, H.E., Gulersonmez, M.C., Stigter, E.C.A., Burgering, B., Geurts, V., Gracanin, A., et al. (2019). Pancreatic cancer organoids recapitulate disease and allow personalized drug screening. Proc. Natl. Acad. Sci. 116, 26580–26590. 10.1073/pnas.1911273116.

36. Gonzalez-Perez, A., Perez-Llamas, C., Deu-Pons, J., Tamborero, D., Schroeder, M.P., Jene-Sanz, A., Santos, A., and Lopez-Bigas, N. (2013). IntOGen-mutations identifies cancer drivers across tumor types. Nat. Methods 10, 1081–1082. 10.1038/nmeth.2642.

37. Belyeu, J.R., Chowdhury, M., Brown, J., Pedersen, B.S., Cormier, M.J., Quinlan, A.R., and Layer, R.M. (2021). Samplot: a platform for structural variant visual validation and automated filtering. Genome Biol. 22, 161. 10.1186/s13059-021-02380-5.

38. Polak, P., Karlić, R., Koren, A., Thurman, R., Sandstrom, R., Lawrence, M.S., Reynolds, A., Rynes, E., Vlahoviček, K., Stamatoyannopoulos, J.A., et al. (2015). Cell-of-origin chromatin organization shapes the mutational landscape of cancer. Nature 518, 360–364. 10.1038/nature14221.

39. Pich, O., Muiños, F., Sabarinathan, R., Reyes-Salazar, I., Gonzalez-Perez, A., and Lopez-Bigas, N. (2018). Somatic and Germline Mutation Periodicity Follow the Orientation of the DNA Minor Groove around Nucleosomes. Cell 175, 1074–1087.e18. 10.1016/j.cell.2018.10.004.

40. Meier, D.T., Rachid, L., Wiedemann, S.J., Traub, S., Trimigliozzi, K., Stawiski, M., Sauteur, L., Winter, D.V., Le Foll, C., Brégère, C., et al. (2022). Prohormone convertase 1/3 deficiency causes obesity due to impaired proinsulin processing. Nat. Commun. 13, 4761. 10.1038/s41467-022-32509-4.

41. Chimienti, F., Devergnas, S., Pattou, F., Schuit, F., Garcia-Cuenca, R., Vandewalle, B., Kerr-Conte, J., Van Lommel, L., Grunwald, D., Favier, A., et al. (2006). In vivo expression and functional characterization of the zinc transporter ZnT8 in glucose-induced insulin secretion. J. Cell Sci. 119, 4199–4206. 10.1242/jcs.03164.

42. Kawase, T., Ichikawa, H., Ohta, T., Nozaki, N., Tashiro, F., Ohki, R., and Taya, Y. (2008). p53 target gene AEN is a nuclear exonuclease required for p53-dependent apoptosis. Oncogene 27, 3797–3810. 10.1038/onc.2008.32.

43. Xie, D., Gore, C., Zhou, J., Pong, R.-C., Zhang, H., Yu, L., Vessella, R.L., Min, W., and Hsieh, J.-T. (2009). DAB2IP coordinates both PI3K-Akt and ASK1 pathways for cell survival and apoptosis. Proc. Natl. Acad. Sci. 106, 19878–19883. 10.1073/pnas.0908458106.

44. Kawase, T., Ohki, R., Shibata, T., Tsutsumi, S., Kamimura, N., Inazawa, J., Ohta, T., Ichikawa, H., Aburatani, H., Tashiro, F., et al. (2009). PH Domain-Only Protein PHLDA3 Is a p53-Regulated Repressor of Akt. Cell 136, 535–550. 10.1016/j.cell.2008.12.002.

45. Lovén, J., Hoke, H.A., Lin, C.Y., Lau, A., Orlando, D.A., Vakoc, C.R., Bradner, J.E., Lee, T.I., and Young, R.A. (2013). Selective Inhibition of Tumor Oncogenes by Disruption of Super-Enhancers. Cell 153, 320–334. 10.1016/j.cell.2013.03.036.

46. Pott, S., and Lieb, J.D. (2015). What are super-enhancers? Nat. Genet. 47, 8–12. 10.1038/ng.3167.

47. Liau, W.S., Tan, S.H., Ngoc, P.C.T., Wang, C.Q., Tergaonkar, V., Feng, H., Gong, Z., Osato, M., Look, A.T., and Sanda, T. (2017). Aberrant activation of the GIMAP enhancer by oncogenic transcription factors in T-cell acute lymphoblastic leukemia. Leukemia 31, 1798–1807. 10.1038/leu.2016.392.

48. Saint-André, V., Federation, A.J., Lin, C.Y., Abraham, B.J., Reddy, J., Lee, T.I., Bradner, J.E., and Young, R.A. (2016). Models of human core transcriptional regulatory circuitries. Genome Res. 26, 385–396. 10.1101/gr.197590.115.

49. Kundaje, A., Meuleman, W., Ernst, J., Bilenky, M., Yen, A., Heravi-Moussavi, A., Kheradpour, P., Zhang, Z., Wang, J., Ziller, M.J., et al. (2015). Integrative analysis of 111 reference human epigenomes. Nature 518, 317–330. 10.1038/nature14248.

50. Lawlor, N., Márquez, E.J., Orchard, P., Narisu, N., Shamim, M.S., Thibodeau, A., Varshney, A., Kursawe, R., Erdos, M.R., Kanke, M., et al. (2019). Multiomic Profiling Identifies cis-Regulatory Networks Underlying Human Pancreatic β Cell Identity and Function. Cell Rep. 26, 788–801.e6. 10.1016/j.celrep.2018.12.083.

51. Bhandare, R., Schug, J., Lay, J.L., Fox, A., Smirnova, O., Liu, C., Naji, A., and Kaestner, K.H. (2010). Genome-wide analysis of histone modifications in human pancreatic islets. Genome Res. 20, 428–433. 10.1101/gr.102038.109.

52. Dunham, I., Kundaje, A., Aldred, S.F., Collins, P.J., Davis, C.A., Doyle, F., Epstein, C.B., Frietze, S., Harrow, J., Kaul, R., et al. (2012). An integrated encyclopedia of DNA elements in the human genome. Nature 489, 57–74. 10.1038/nature11247.

53. Bramswig, N.C., Everett, L.J., Schug, J., Dorrell, C., Liu, C., Luo, Y., Streeter, P.R., Naji, A., Grompe, M., and Kaestner, K.H. (2013). Epigenomic plasticity enables human pancreatic α to β cell reprogramming. J. Clin. Invest. 123, 1275–1284. 10.1172/JCI66514.

54. Thomsen, S.K., and Gloyn, A.L. (2014). The pancreatic β cell: recent insights from human genetics. Trends Endocrinol. Metab. 25, 425–434. 10.1016/j.tem.2014.05.001.

55. Eizirik, D.L., Pasquali, L., and Cnop, M. (2020). Pancreatic β-cells in type 1 and type 2 diabetes mellitus: different pathways to failure. Nat. Rev. Endocrinol. 16, 349–362. 10.1038/s41574-020-0355-7.

56. Nakagawa, H., and Fujita, M. (2018). Whole genome sequencing analysis for cancer genomics and precision medicine. Cancer Sci. 109, 513–522. 10.1111/cas.13505.

57. Mukherjee, S., Chaturvedi, P., Rankin, S.A., Fish, M.B., Wlizla, M., Paraiso, K.D., MacDonald, M., Chen, X., Weirauch, M.T., Blitz, I.L., et al. (2020). Sox17 and β-catenin co-occupy Wnt-responsive enhancers to govern the endoderm gene regulatory network. eLife 9, e58029. 10.7554/eLife.58029.

58. Collinson, A., Collier, A.J., Morgan, N.P., Sienerth, A.R., Chandra, T., Andrews, S., and Rugg-Gunn, P.J. (2016). Deletion of the Polycomb-Group Protein EZH2 Leads to Compromised Self-Renewal and Differentiation Defects in Human Embryonic Stem Cells. Cell Rep. 17, 2700–2714. 10.1016/j.celrep.2016.11.032.

59. Krücken, J., Schroetel, R.M.U., Müller, I.U., Saïdani, N., Marinovski, P., Benten, W.P.M., Stamm, O., and Wunderlich, F. (2004). Comparative analysis of the human gimap gene cluster encoding a novel GTPase family. Gene 341, 291–304. 10.1016/j.gene.2004.07.005.

60. Schmidl, C., Rendeiro, A.F., Sheffield, N.C., and Bock, C. (2015). ChIPmentation: fast, robust, low-input ChIP-seq for histones and transcription factors. Nat. Methods 12, 963–965. 10.1038/nmeth.3542.

61. Kaya-Okur, H.S., Wu, S.J., Codomo, C.A., Pledger, E.S., Bryson, T.D., Henikoff, J.G., Ahmad, K., and Henikoff, S. (2019). CUT&Tag for efficient epigenomic profiling of small samples and single cells. Nat. Commun. 10, 1930. 10.1038/s41467-019-09982-5.

62. Patro, R., Duggal, G., Love, M.I., Irizarry, R.A., and Kingsford, C. (2017). Salmon provides fast and bias-aware quantification of transcript expression. Nat. Methods 14, 417–419. 10.1038/nmeth.4197.

63. Soneson, C., Love, M.I., and Robinson, M.D. (2015). Differential analyses for RNA-seq: transcript-level estimates improve gene-level inferences. F1000Research 4, 1521. 10.12688/f1000research.7563.1.

64. Langmead, B., and Salzberg, S.L. (2012). Fast gapped-read alignment with Bowtie 2. Nat. Methods 9, 357–359. 10.1038/nmeth.1923.

65. Li, H., Handsaker, B., Wysoker, A., Fennell, T., Ruan, J., Homer, N., Marth, G., Abecasis, G., Durbin, R., and 1000 Genome Project Data Processing Subgroup (2009). The Sequence Alignment/Map format and SAMtools. Bioinformatics 25, 2078–2079. 10.1093/bioinformatics/btp352.

66. Ross-Innes, C.S., Stark, R., Teschendorff, A.E., Holmes, K.A., Ali, H.R., Dunning, M.J., Brown, G.D., Gojis, O., Ellis, I.O., Green, A.R., et al. (2012). Differential oestrogen receptor binding is associated with clinical outcome in breast cancer. Nature 481, 389–393. 10.1038/nature10730.

67. Liao, Y., Smyth, G.K., and Shi, W. (2019). The R package Rsubread is easier, faster, cheaper and better for alignment and quantification of RNA sequencing reads. Nucleic Acids Res. 47, e47. 10.1093/nar/gkz114.

68. Love, M.I., Huber, W., and Anders, S. (2014). Moderated estimation of fold change and dispersion for RNA-seq data with DESeq2. Genome Biol. 15, 550. 10.1186/s13059-014-0550-8.

69. Fishilevich, S., Nudel, R., Rappaport, N., Hadar, R., Plaschkes, I., Iny Stein, T., Rosen, N., Kohn, A., Twik, M., Safran, M., et al. (2017). GeneHancer: genome-wide integration of enhancers and target genes in GeneCards. Database 2017. 10.1093/database/bax028.

70. Gu, Z., and Hübschmann, D. (2023). *rGREAT* : an R/bioconductor package for functional enrichment on genomic regions. Bioinformatics 39, btac745. 10.1093/bioinformatics/btac745.

71. Australian Pancreatic Cancer Genome Initiative, Scarpa, A., Chang, D.K., Nones, K., Corbo, V., Patch, A.-M., Bailey, P., Lawlor, R.T., Johns, A.L., Miller, D.K., et al. (2017). Whole-genome landscape of pancreatic neuroendocrine tumours. Nature 543, 65–71. 10.1038/nature21063.

72. Wang, H., Bender, A., Wang, P., Karakose, E., Inabnet, W.B., Libutti, S.K., Arnold, A., Lambertini, L., Stang, M., Chen, H., et al. (2017). Insights into beta cell regeneration for diabetes via integration of molecular landscapes in human insulinomas. Nat. Commun. 8, 767. 10.1038/s41467-017-00992-9.

73. Cao, Y., Gao, Z., Li, L., Jiang, X., Shan, A., Cai, J., Peng, Y., Li, Y., Jiang, X., Huang, X., et al. (2013). Whole exome sequencing of insulinoma reveals recurrent T372R mutations in YY1. Nat. Commun. 4, 2810. 10.1038/ncomms3810.

74. Kim, S., Scheffler, K., Halpern, A.L., Bekritsky, M.A., Noh, E., Källberg, M., Chen, X., Kim, Y., Beyter, D., Krusche, P., et al. (2018). Strelka2: fast and accurate calling of germline and somatic variants. Nat. Methods 15, 591–594. 10.1038/s41592-018-0051-x.

75. Benjamin, D., Sato, T., Cibulskis, K., Getz, G., Stewart, C., and Lichtenstein, L. (2019). Calling Somatic SNVs and Indels with Mutect2 (Bioinformatics) 10.1101/861054.

76. Chen, X., Schulz-Trieglaff, O., Shaw, R., Barnes, B., Schlesinger, F., Källberg, M., Cox, A.J., Kruglyak, S., and Saunders, C.T. (2016). Manta: rapid detection of structural variants and indels for germline and cancer sequencing applications. Bioinformatics 32, 1220–1222. 10.1093/bioinformatics/btv710.

77. Chen, S., Francioli, L.C., Goodrich, J.K., Collins, R.L., Kanai, M., Wang, Q., Alföldi, J., Watts, N.A., Vittal, C., Gauthier, L.D., et al. (2022). A genome-wide mutational constraint map quantified from variation in 76,156 human genomes (Genetics) 10.1101/2022.03.20.485034.

78. Sherry, S.T. (2001). dbSNP: the NCBI database of genetic variation. Nucleic Acids Res. 29, 308–311. 10.1093/nar/29.1.308.

79. Wang, K., Li, M., and Hakonarson, H. (2010). ANNOVAR: functional annotation of genetic variants from high-throughput sequencing data. Nucleic Acids Res. 38, e164–e164. 10.1093/nar/gkq603.

80. Mayakonda, A., Lin, D.-C., Assenov, Y., Plass, C., and Koeffler, H.P. (2018). Maftools: efficient and comprehensive analysis of somatic variants in cancer. Genome Res. 28, 1747–1756. 10.1101/gr.239244.118.

81. Alexandrov, L.B., Kim, J., Haradhvala, N.J., Huang, M.N., Tian Ng, A.W., Wu, Y., Boot, A., Covington, K.R., Gordenin, D.A., Bergstrom, E.N., et al. (2020). The repertoire of mutational signatures in human cancer. Nature 578, 94–101. 10.1038/s41586-020-1943-3.

82. Rausch, T., Zichner, T., Schlattl, A., Stütz, A.M., Benes, V., and Korbel, J.O. (2012). DELLY: structural variant discovery by integrated paired-end and split-read analysis. Bioinformatics 28, i333–i339. 10.1093/bioinformatics/bts378.

83. Cameron, D.L., Baber, J., Shale, C., Valle-Inclan, J.E., Besselink, N., Van Hoeck, A., Janssen, R., Cuppen, E., Priestley, P., and Papenfuss, A.T. (2021). GRIDSS2: comprehensive characterisation of somatic structural variation using single breakend variants and structural variant phasing. Genome Biol. 22, 202. 10.1186/s13059-021-02423-x.

84. Pedersen, B.S., and Quinlan, A.R. (2019). Duphold: scalable, depth-based annotation and curation of high-confidence structural variant calls. GigaScience 8. 10.1093/gigascience/giz040.

85. Chowdhury, M., Pedersen, B.S., Sedlazeck, F.J., Quinlan, A.R., and Layer, R.M. (2022). Searching thousands of genomes to classify somatic and novel structural variants using STIX. Nat. Methods 19, 445–448. 10.1038/s41592-022-01423-4.

86. Belyeu, J.R., Chowdhury, M., Brown, J., Pedersen, B.S., Cormier, M.J., Quinlan, A.R., and Layer, R.M. (2021). Samplot: a platform for structural variant visual validation and automated filtering. Genome Biol. 22, 161. 10.1186/s13059-021-02380-5.

87. Harrow, J., Frankish, A., Gonzalez, J.M., Tapanari, E., Diekhans, M., Kokocinski, F., Aken, B.L., Barrell, D., Zadissa, A., Searle, S., et al. (2012). GENCODE: The reference human genome annotation for The ENCODE Project. Genome Res. 22, 1760–1774. 10.1101/gr.135350.111.

88. Gel, B., Díez-Villanueva, A., Serra, E., Buschbeck, M., Peinado, M.A., and Malinverni, R. (2016). regioneR: an R/Bioconductor package for the association analysis of genomic regions based on permutation tests. Bioinformatics 32, 289–291. 10.1093/bioinformatics/btv562.

89. Martin Morgan and Hervé Pagès and Valerie Obenchain and Nathaniel Hayden (2022). Rsamtools: Binary alignment (BAM), FASTA, variant call (BCF), and tabix file import.

90. Siepel, A., Bejerano, G., Pedersen, J.S., Hinrichs, A.S., Hou, M., Rosenbloom, K., Clawson, H., Spieth, J., Hillier, L.W., Richards, S., et al. (2005). Evolutionarily conserved elements in vertebrate, insect, worm, and yeast genomes. Genome Res. 15, 1034–1050. 10.1101/gr.3715005.

91. Hadley Wickham (2016). ggplot2: Elegant Graphics for Data Analysis (Springer-Verlag New York).

92. Korotkevich, G., Sukhov, V., Budin, N., Shpak, B., Artyomov, M.N., and Sergushichev, A. (2016). Fast gene set enrichment analysis (Bioinformatics) 10.1101/060012.

93. Wu, T., Hu, E., Xu, S., Chen, M., Guo, P., Dai, Z., Feng, T., Zhou, L., Tang, W., Zhan, L., et al. (2021). clusterProfiler 4.0: A universal enrichment tool for interpreting omics data. The Innovation 2, 100141. 10.1016/j.xinn.2021.100141.

94. Guangchuang Yu (2018). enrichplot. (Bioconductor). 10.18129/B9.BIOC.ENRICHPLOT 10.18129/B9.BIOC.ENRICHPLOT.

95. Heinz, S., Benner, C., Spann, N., Bertolino, E., Lin, Y.C., Laslo, P., Cheng, J.X., Murre, C., Singh, H., and Glass, C.K. (2010). Simple Combinations of Lineage-Determining Transcription Factors Prime cis-Regulatory Elements Required for Macrophage and B Cell Identities. Mol. Cell 38, 576–589. 10.1016/j.molcel.2010.05.004.

96. Coetzee, S.G., Coetzee, G.A., and Hazelett, D.J. (2015). *motifbreakR* : an R/Bioconductor package for predicting variant effects at transcription factor binding sites. Bioinformatics 31, 3847–3849. 10.1093/bioinformatics/btv470.

97. Paul Shannon and Matt Richards (2022). MotifDb: An Annotated Collection of Protein-DNA Binding Sequence Motifs.

98. Lawrence, M., Gentleman, R., and Carey, V. (2009). rtracklayer: an R package for interfacing with genome browsers. Bioinformatics 25, 1841–1842. 10.1093/bioinformatics/btp328.

