## Supplementary Materials for "Implications of noncoding regulatory functions in the development of insulinomas"

### Supplementary Tables:

**Table S1. Description of insulinoma samples.** Columns "RNA-seq", "H3K27ac", "WGS" and "WES" indicate if the experiment was performed in that sample ("Y") or not ("N"). "Orignal ID" contains the identifier used in the original publication.

**Table S2**. **Differentially expressed genes**. List of differentially expressed genes when comparing insulinomas with human islets (Comparison = INSvsHI) or FACS-purified β cells (Comparison = INSvsBETA).

**Table S3. Consensus regulatory elements**. Stringent consensus H3K27ac regions, including differential state when comparing insulinomas to human islets ("type"), wheter the region is located in an IRD ("IRD") and if it presents the H3K27me3 repressive mark in control tissues (human islets or EndoC-βH1, "H3K27me3 in controls").

**Table S4**. **Mutations of insulinoma cohort**. List of mutations along the insulinoma cohort including coding, non-coding and structural variants.

**Table S5. Variant Regulatory Elements**. H3K27ac sites bearing an insulinoma somatic mutation. The column "Associated Gene" contains genes 1) whose TSS is closer than 5 kb upstream and 1 kb downstream and 2) nearest upstream and downstream gene TSS within a 1 Mb distance.

**Table S6. Insulinoma Regulatory Domains**. Coordinates of IRDs, including the chromosome-specific distance cutoff used to include REs in the domain ("Distance cutoff"), the number of RE sites included ("Number of REs") and the peak identifiers of these Res ("RE peak IDs").

**Table S7. Published datasets used in the paper**. List of all published datasets employed in this publication.

### Supplementary Figures:

**Figure S1**. RNA-seq datasets.

**Figure S2**. H3K27ac datasets.

**Figure S3**. H3K27ac signal as a measure of tumor sample heterogeneity.

**Figure S4**. Genetic profile of insulinomas.

**Figure S5**. Non-coding genetic alterations in insulinomas

**Figure S6**. Insulinoma Regulatory Domains (IRDs) specifications.

**Figure S7**. IRDs are polycomb-repressed in untransformed cell types.

#### Figure S1

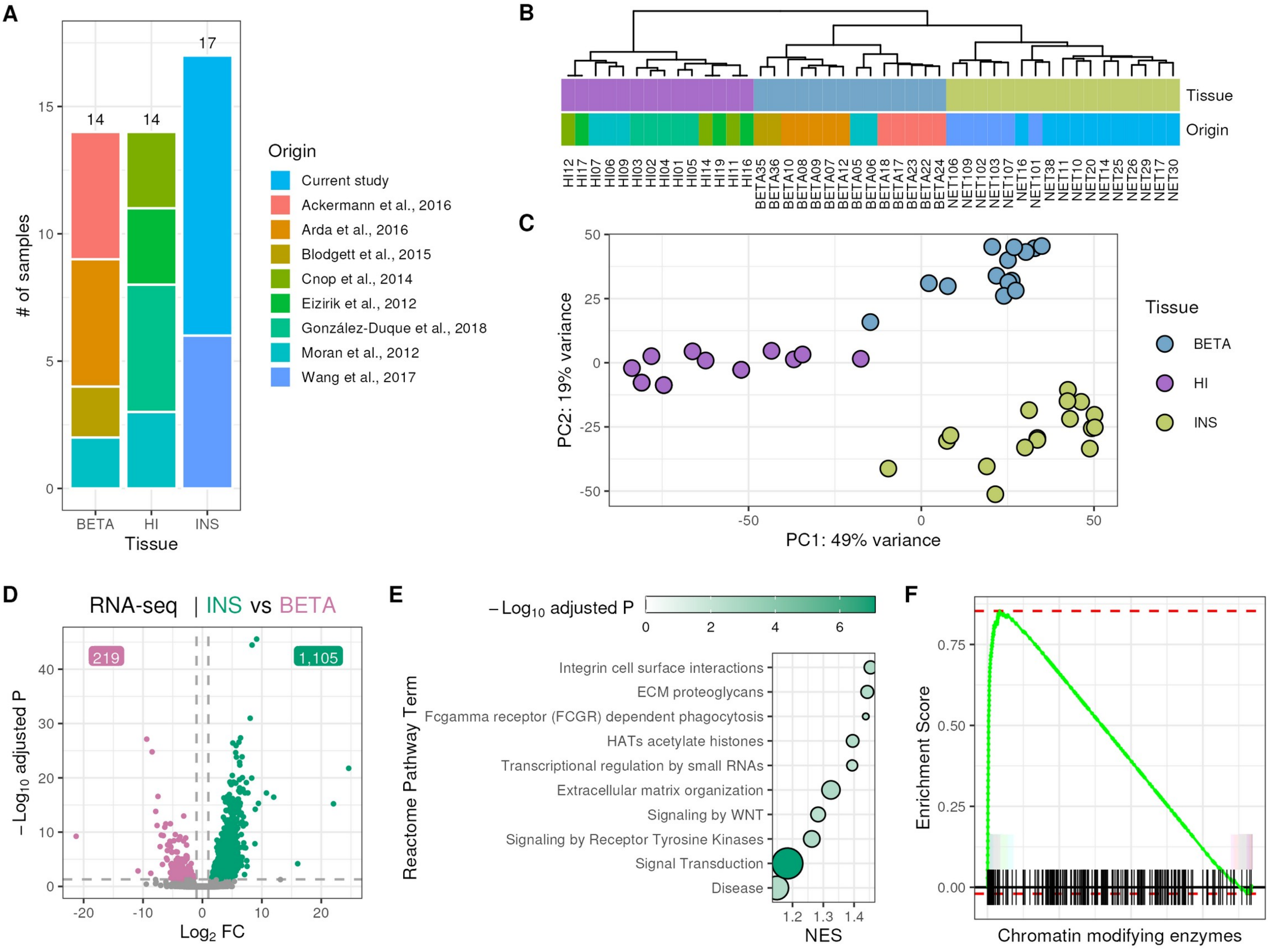


**Figure S1: RNA-seq datasets**. **A**, Origin of the RNA-seq samples used in this project. "Current study" indicates the data generated in this publication. **B**, Unsupervised hierarchical clustering of samples using expression of protein-coding genes obtained from RNA-seq experiments. **C**, Principal component analysis of the 1,000 most variably expressed genes. For **B** and **C**, data is transformed using the variance stabilizing transformation (VST — see Methods). **D**, Volcano plot of differentially expressed genes in insulinomas vs β cells. Dotted lines show thresholds for significance (|$\log_2$ fold change| > 1 and adjusted P < 0.05). **E**, GSEA of Reactome Pathway terms positively enriched in insulinomas compared to β cells. **F**, GSEA of the term "HATs acetylate histones" in up-regulated genes in insulinoma compared to human islets.

#### Figure S2

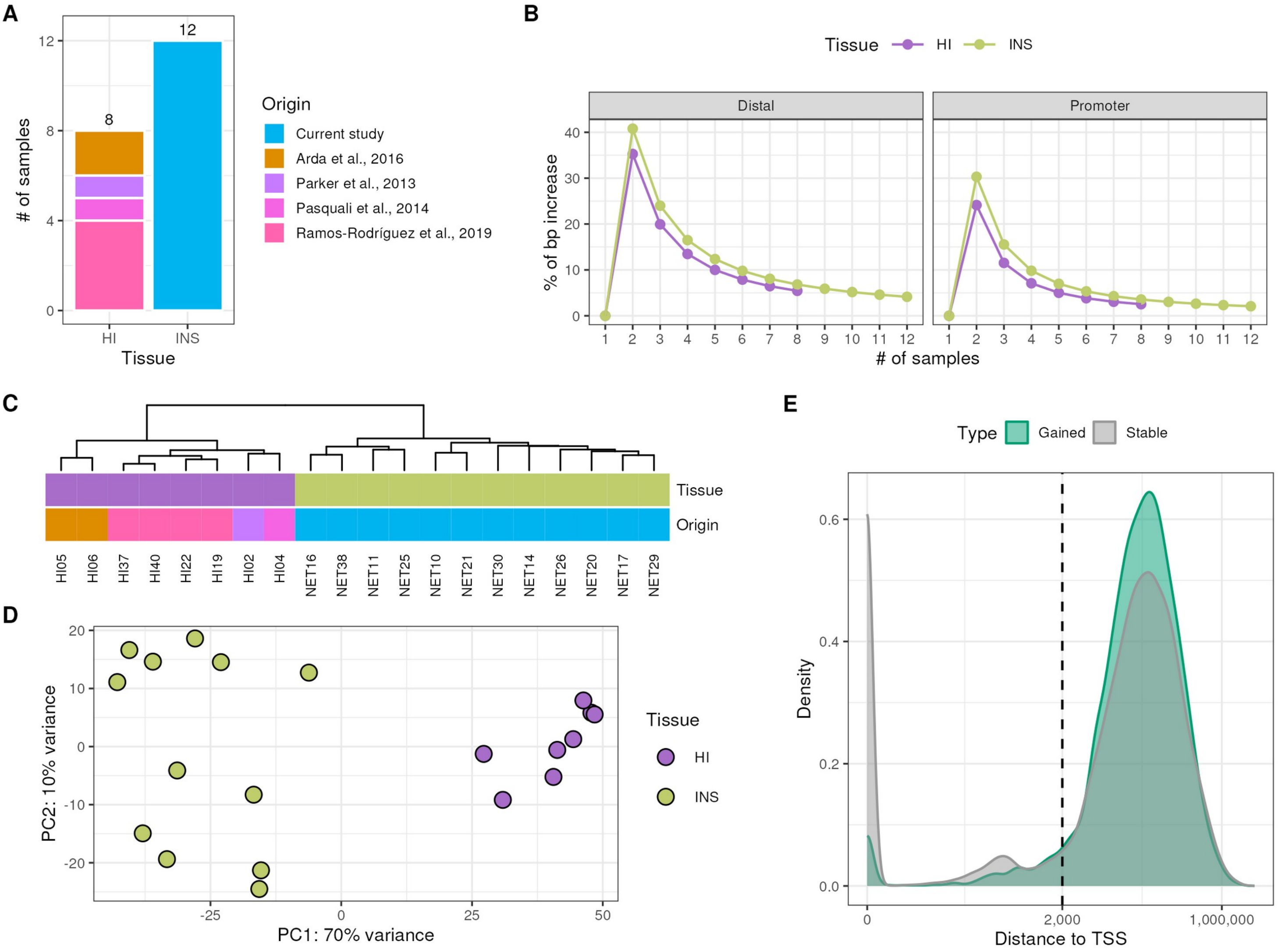


**Figure S2: H3K27ac datasets**. **A**, Origin of the H3K27ac ChIP-seq samples used in this project. "Current study" indicates the data generated in this publication. **B**, Saturation plots indicating the additional base pairs covered by H3K27ac peaks obtained by combining an increasing number of samples (HI: Human Islets; INS: insulinomas). The dot represents the average base-pair increase derived by permutation of all different sample combinations. The number of permutated samples is indicated in the x axis. **C**, Unsupervised hierarchical clustering of samples based on H3K27ac enriched sites. **D**, Principal component analysis of the 1,000 most variable H3K27ac regions. For **C** and **D,** data is transformed using the variance stabilizing transformation (VST — see Methods). **E**, Distribution of distances to the nearest protein-coding TSS for H3K27ac enriched sites.

#### Figure S3

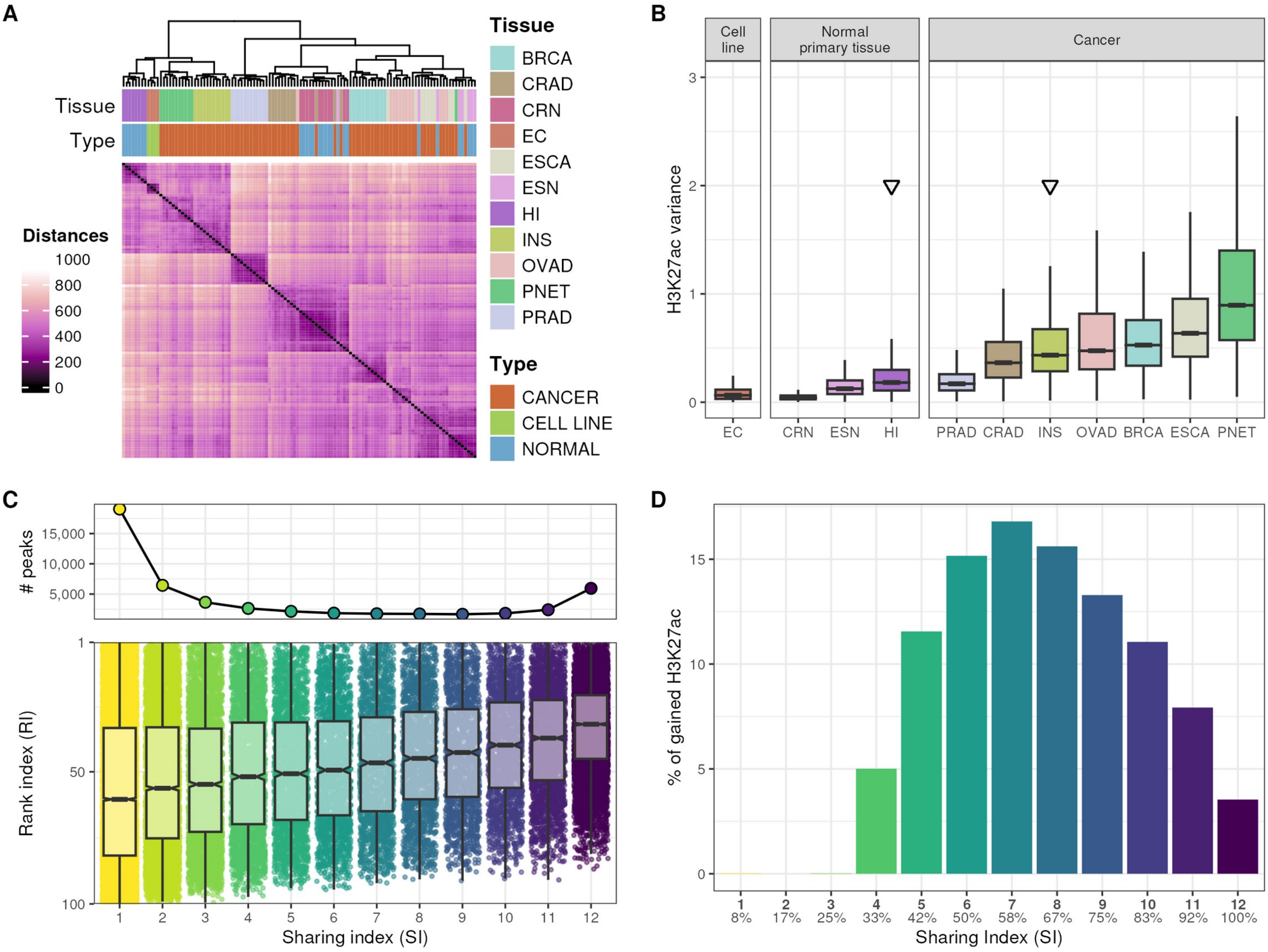

A
Tissue
Type
Distances
1000
800
600
400
200
0
Tissue
BRCA
CRAD
CRN
EC
ESCA
ESN
HI
INS
OVAD
PNET
PRAD
Type
CANCER
CELL LINE
NORMAL
B
Cell line
Normal primary tissue
Cancer
H3K27ac variance
EC
CRN
ESN
HI
PRAD
CRAD
INS
OVAD
BRCA
ESCA
PNET
C
### peaks
15,000
10,000
5,000
Rank index (RI)
Sharing index (SI)
D
% of gained H3K27ac
Sharing Index (SI)
8%
17%
25%
33%
42%
50%
58%
67%
75%
83%
92%
100%

#### Figure S4

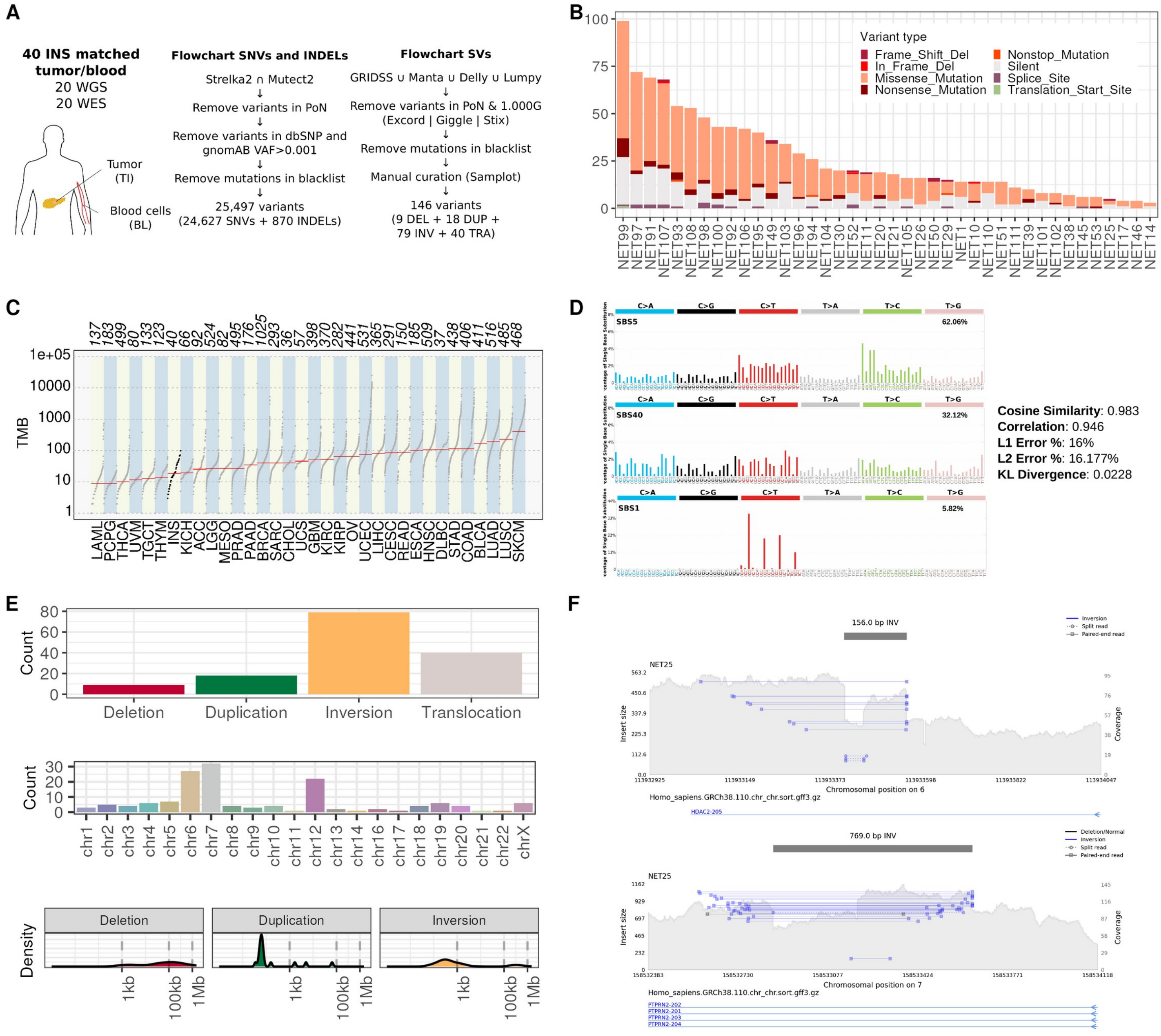


**Figure S4: Genetic profile of insulinomas. A,** Schematic representation of the algorithms used for calling SNVs, INDELs and SVs in this study, indicating the final number of mutations obtained after filtering and manual curation. **B**, Distribution of coding mutations across the insulinoma cohort. Each bar represents the number and type of coding mutations per sample. **C**, The tumor mutation burden of insulinomas (black bold dots) and that of 33 different tumor types (TCGA data) is represented. Each dot represents a tumor sample, while an horizontal line depicts the median mutation load for each cancer type. **D**, The distinct SBS *de novo* signatures observed in insulinomas are decomposed and matched to COSMIC annotations. The cosine similarity, correlation, L1 error %, L2 error %, and KL divergence are reported. The mutation signature profile of insulinomas matches those related to aging and clock-like signatures (SBS1, SBS5 and SBS40). **E**, Distribution of curated insulinoma SVs in terms of their type (top), genome positioning (middle) and size (bottom). **F**, Samplot of manually curated SV found at the *HDAC2* and *PTPRN2* loci.

#### Figure S5

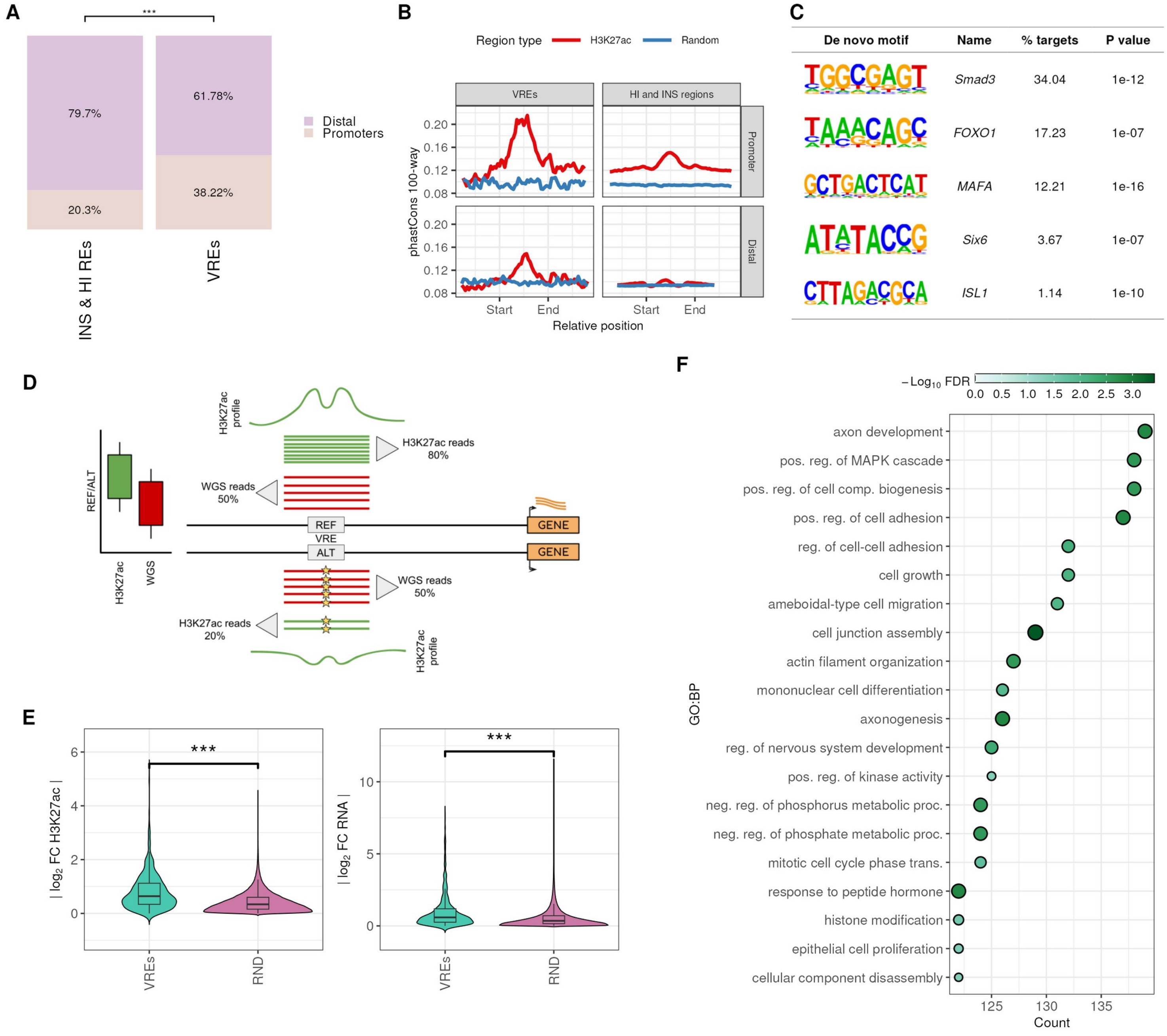


**Figure S5: Non-coding genetic alterations in insulinomas. A**, Genomic localization of VREs compared to non-mutated H3K27ac sites active in insulinoma and human islet samples. Regions within 2kb of the gene TSS are categorized as promoters, while those beyond this range are annotated distal. $\chi^2$ test ***P < 0.001 **B**, Mean sequence conservation scores at H3K27ac sites bearing (VRE) or not a somatic mutation annotated according to their genomic localization (Distal, Promoter). Peaks were extended from the center 1 kb to each direction and mean score was calculated in 20 bp windows. **C**, Predominant *de novo* motifs identified by HOMER in VREs (score > 0.7 and P < $1x10^{-7}$). **D**, Representative scheme of the methodology used to compute differential enrichment of the H3K27ac histone modification at VREs depending on the somatic mutation genotype (see **Figure 2D**). **E**, Violin plots showing the distribution of H3K27ac (left) and RNA-seq (right) absolute fold changes, when comparing mutated against wildtype VREs. In the matched control, the genotype is randomly assigned (RND). Two-sided Wilcoxon test ***P < 0.001. **F**, Gene Ontology Biological Process annotation for genes associated with all genomic aberrations detected in the insulinoma cohort.

#### Figure S6

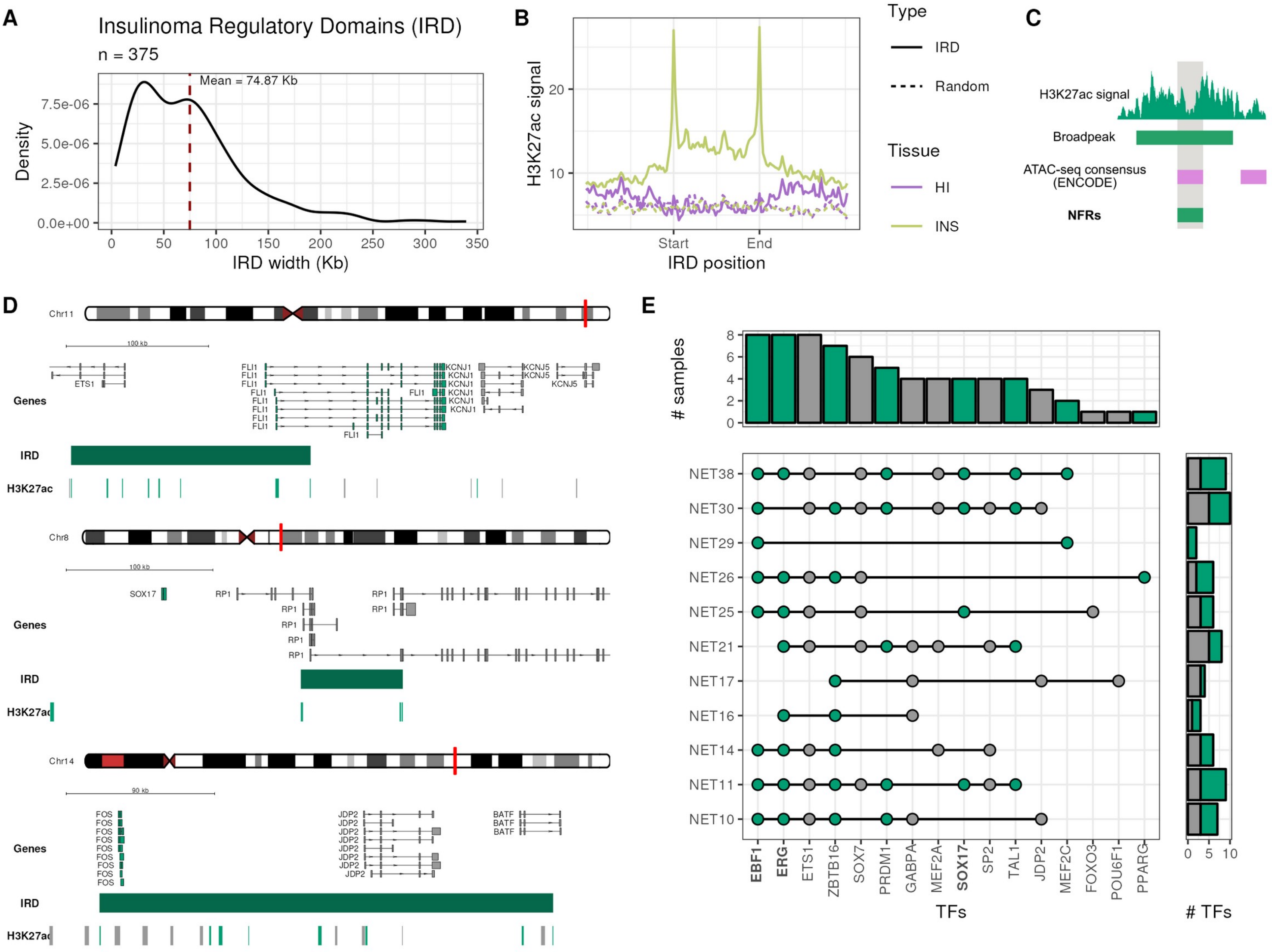


**Figure S6: Insulinoma Regulatory Domains (IRDs) specifications**. **A**, Distribution of Insulinoma Regulatory Domain (IRD) widths. Chromosome-specific thresholds for stitching together REs into an IRD range from 4.3 to 126.8 Kb (see **Table S6**). **B**, Average H3K27ac signal at IRD in human islets (HI), insulinomas (INS) and their matched randomized sets. **C**, Scheme illustrating the methodology used to detect nucleosome free regions (NFRs) used to derive binding motifs (see **Figure 3E**). **D**, Examples of TFs identified from *de novo* motif analysis in IRDs, that are themselves regulated by an IRD, thus matching the definition of Core transcriptional Regulatory Circuitries (CRC). **E**, Top CRC identified in each insulinoma sample. Fill represents TF gene type (stable: grey; up-regulated: green; down-regulated: pink). Names in bold correspond to TFs that were also identified in the *de novo* motif analysis.

### Figure S7

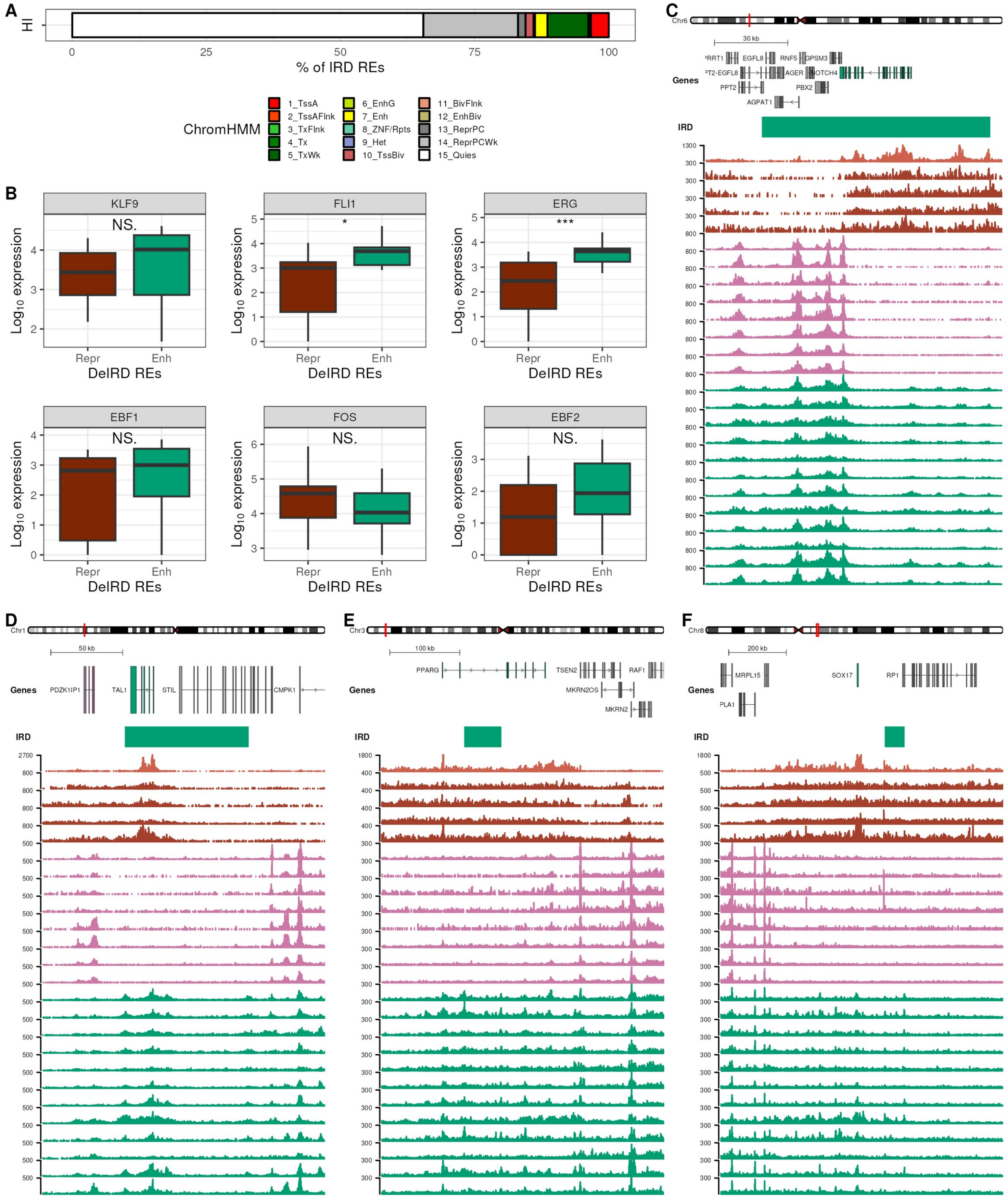

A
HI
0
25
50
75
100
% of IRD REs
ChromHMM
1_TssA
2_TssAFlnk
3_TxFlnk
4_Tx
5_TxWk
6_EnhG
7_Enh
8_ZNF/Rpts
9_Het
10_TssBiv
11_BivFlnk
12_EnhBiv
13_ReprPC
14_ReprPCWk
15_Quies
B
KLF9
NS.
FLI1
*
ERG
***
EBF1
NS.
FOS
NS.
EBF2
NS.
Log10 expression
Repr
Enh
DeIRD REs
C
Chr6
30 kb
Genes
EGFL8
RNF5
GPSM3
AGER
NOTCH4
PPT2
PBX2
AGPAT1
IRD
D
Chr1
50 kb
Genes
PDZK1IP1
TAL1
STIL
CMPK1
IRD
E
Chr3
100 kb
Genes
PPARG
TSEN2
RAF1
MKRN2OS
MKRN2
IRD
F
Chr8
200 kb
Genes
MRPL15
SOX17
RP1
IRD

**Figure S7: IRDs are polycomb-repressed in untransformed cell types**. **A**, Chromatin state annotation of IRDs based on the Epigenome Roadmap ChromHMM computed in human pancreatic islets. **B**, Expression of TFs acting at IRD (**Figure 3E**) in Epigenome Roadmap tissues in which DeIRDs are preferentially annotated as "polycomb-repressed" (*Repr*, $\log_2$ ratio < -0.58) or as "enhancers" (*Enh*, $\log_2$ ratio > 0.58). Two-sided Wilcoxon test. * P < 0.05; *** P < 0.001; NS: Not Significant. **C-F**, The *NOTCH4* (**C**), *TAL1* (**D**), *PPARG* (**E**) and *SOX17* (**F**) loci as representative examples of DeIRD genes whose activation may be associated with tumor development. Red: H3K27me3 EndoC-βH1 (light) and human islets (dark); Pink: H3K27ac in human islets; Green: H3K27ac in insulinomas.
